# Rapid removal of nuclear aggregates via proteasome- and VCP-dependent disaggregation

**DOI:** 10.64898/2026.07.20.739537

**Authors:** Giel Korsten, Grace P. Smith, Wilco Nijenhuis, Anne F. J. Janssen, Lukas C. Kapitein

## Abstract

Cells use multiple protein quality control (PQC) mechanisms to counteract the toxic effects of protein aggregation caused by cellular stress, ageing or disease. While it is known that PQC mechanisms differ between cellular compartments, studying such differences has remained challenging. Previously, we developed an assay to study cytosolic PQC using aggregates formed through chemically-induced dimerization (termed PIMs, particles induced by multimerization). Here, we introduce nuclear PIMs as a tool to study nuclear quality control. Using high-resolution and high-throughput imaging, we show that nuclear aggregate removal depends on the proteasome and the unfoldase VCP, but not on Hsp70. Strikingly, following dissolution many PIM subunits were exported to the cytosol via exportin-1-dependent shuttling, indicating that disaggregation and resolubilization dominated over degradation. Proteasomal disaggregation was confirmed using live-cell turnover experiments. Together, these findings reveal a mechanism in which the proteasome and VCP disaggregate, rather than degrade, nuclear protein aggregates, with the resulting subunits subsequently cleared via cytosolic aggrephagy.

## Introduction

Accumulation of protein aggregates in cells is a well-known feature of various diseases, including both neurodegenerative and ageing-related conditions (Wilson et al., 2023). To prevent protein aggregation, a complex protein quality control (PQC) network functions to recognize unfolded proteins that are prone to aggregate and to facilitate refolding or degradation (Nillegoda et al., 2018; Rosenzweig et al., 2019). However, these preventative mechanisms can be overwhelmed by cell stress or the prolonged presence of mutant proteins, or become impaired due to aging or disease (Hipp et al., 2019). The resulting protein aggregates can be harmful to cells and result in downstream effects including toxic gain- or loss of function (Wilson et al., 2023). Therefore, cells have specialized mechanisms that facilitate removal of protein aggregates.

In the cytosol, protein aggregates can be delivered to the lysosome as intact aggregates in a process called autophagy (Lamark & Johansen, 2012). Although autophagy has been implicated in some aspects of nuclear PQC, such as removal of yeast nuclear pore complexes and mammalian lamins (Dou et al., 2015; C. W. Lee et al., 2020; Z. Li & Nakatogawa, 2022), its contribution to nuclear aggregate clearance is likely limited. Therefore, other mechanisms are needed for effective PQC of nuclear aggregates (Iwata et al., 2005). In yeast, early work identified disaggregation activity of the heat shock protein Hsp70, together with the AAA-ATPase Hsp104 (Glover & Lindquist, 1998; Mogk et al., 2018; Parsell et al., 1994; Weibezahn et al., 2004). Metazoans lack an Hsp104 homologue and rely on a complex set of co-chaperones to enable disaggregation. Here, Hsp70 acts together with nucleotide exchange factors (NEFs) of the Hsp110 family, and Hsp40s, a highly diverse class of J-domain containing proteins (Kampinga & Craig, 2010; Nillegoda et al., 2015). This triad functions in the nucleus and cytosol, with a varying composition of players and isoforms of Hsp70, NEFs and J-domain proteins likely regulating client recognition, activity, and substrate fate in each compartment (den Brave et al., 2020; Hjerpe et al., 2016; Kampinga & Craig, 2010; Serlidaki et al., 2020). However, the metazoan Hsp70 triad is generally deemed a less efficient disaggregase compared to the yeast system (Duennwald et al., 2012; Shorter, 2011), and our understanding of its role in nuclear disaggregation is limited.

Another major component of the PQC system active in the nucleus is the proteasome. The 26S proteasome is a large multisubunit protein complex. It consists of a cylindrical 20S core particle (CP) that is catalytically active, and one or two 19S regulatory particles (RPs) that are involved in substrate recognition and unfolding, and associate with the 20S barrel at each end (Tomko and Hochstrasser 2013). Fully assembled and functional proteasomes are present in the nucleus, and abundant evidence suggests their involvement in nuclear PQC of a wide range of clients (Albert et al., 2017; Chen & Madura, 2014; Chowdhury & Enenkel, 2015; Enam et al., 2018; Jones & Gardner, 2016; S. H. Park et al., 2013; Prasad et al., 2010, 2018; Shakya et al., 2021). While proteasomes are primarily recognized as proteases that can degrade solubilized clients, for example after solubilization by the Hsp70-triad (Hjerpe et al., 2016), in vitro assays show they can also display disaggregase activity (Cliffe et al., 2019). Moreover, recent work identified the 19S subunit as a part of a cytosolic fragmentase machinery needed for subsequent aggrephagy (Mauthe et al., 2025). Indeed, part of the 19S RP mechanistically and structurally resembles classical AAA-ATPase disaggregases such as yeast Hsp104 (Heuck et al., 2016). Specifically, in the RP, AAA-ATPase subunits organize as a heterohexametric ring and facilitate repeated ATPase dependent conformational changes thought to be required for substrate unfolding (Śledź et al., 2013; Zhang et al., 2009; Zhu et al., 2018). Nonetheless, whether this independent action of the proteasome represents a general mechanism for aggregate removal within cells remains unknown.

Lastly, nuclear disaggregation of protein aggregates might be facilitated by other PQC players. These could include alternative AAA-ATPase proteins present in mammals, such the ubiquitously expressed VCP/p97 (VCP hereafter). VCP is involved in a myriad of PQC processes, including protein complex disassembly, ERAD (ER associated protein degradation), UPS (Ubiquitin Proteasome System) substrate processing, and autophagy (Gallagher et al., 2014; Hill et al., 2021; Meyer et al., 2012; Meyer & Weihl, 2014; Wrobel et al., 2024). More importantly, recent work has shown that VCP can disaggregate cytosolic tau and huntingtin fibrils (Benn et al., 2024; Ghosh et al., 2018; Saha et al., 2023), and reduce the formation of intranuclear TDP-43 inclusions (Phan et al., 2024).

While the components described above are all implicated in aspects of nuclear PQC, a unified understanding of aggregate removal from the nucleus is lacking. Furthermore, investigating nuclear PQC of protein aggregates has been challenging since it has relied on *in vitro* reconstitutions, induction of protein stress or overexpression of potentially toxic, disease related proteins (Phan et al., 2024; Saha et al., 2023). The latter approaches likely cause off target effects that could impact PQC indirectly, while *in vitro* experimentation fails to accurately mimic the complexity of a multicompartment system.

Here, we study nuclear aggregate clearance using a tool that enables the generation of inducible nuclear protein aggregates in cells without the need of toxic stressors. By combining this tool with high throughput live- and fixed-cell imaging, we reveal that nuclear aggregate clearance occurs through disaggregation, followed by nuclear export and subsequent cytosolic reaggregation. This mechanism depends on the proteasome and VCP, but not on the classical Hsp70 chaperone. Live-cell and biochemical analysis of the dynamics of PIM clearance revealed that it is a fast (∼ 5 hours) multistep disaggregation process that prioritizes nuclear export over proteolysis, and leads to cytosolic reaggregation and lysosomal targeting via aggrephagy. Together, these results demonstrate a novel pathway for nuclear aggregate PQC and provide a versatile basis for future high-throughput screening approaches.

## Results

### PIM-mCh-NLS as a tool to study PQC of nuclear protein aggregates

We previously introduced particles induced by multimerization (PIMs) as an inducible tool to study autophagy of protein aggregates (aggrephagy) (Janssen et al., 2018, 2021). PIM monomers are comprised of multiple homodimerization domains that cluster upon the addition of rapalog2. The addition of an EGFP-mCherry dual tag, or the pH sensitive fluorophore mKeima, then enables direct observation of aggregate acidification via aggrephagy. While induction of dualPIM aggregates in cells primarily resulted in the formation of cytosolic aggregates, a small fraction of aggregates was also formed inside the nucleus. Interestingly, these nuclear PIMs were almost completely cleared from the nucleus within 1-2 hours (Fig. 1, A). In order to selectively study nuclear aggregate removal, we generated a mCherry tagged PIM construct carrying a c-terminal nuclear localization signal (NLS; Fig 1, B). To enable robust, high throughput analysis of nuclear aggregate clearance we integrated this construct into RPE1-FRT/TR cells, generating a stable cell line that expresses PIM-mCh-NLS under a doxycycline inducible promotor (hereafter called RPE-PIM-mCh-NLS).

**Figure 1:**
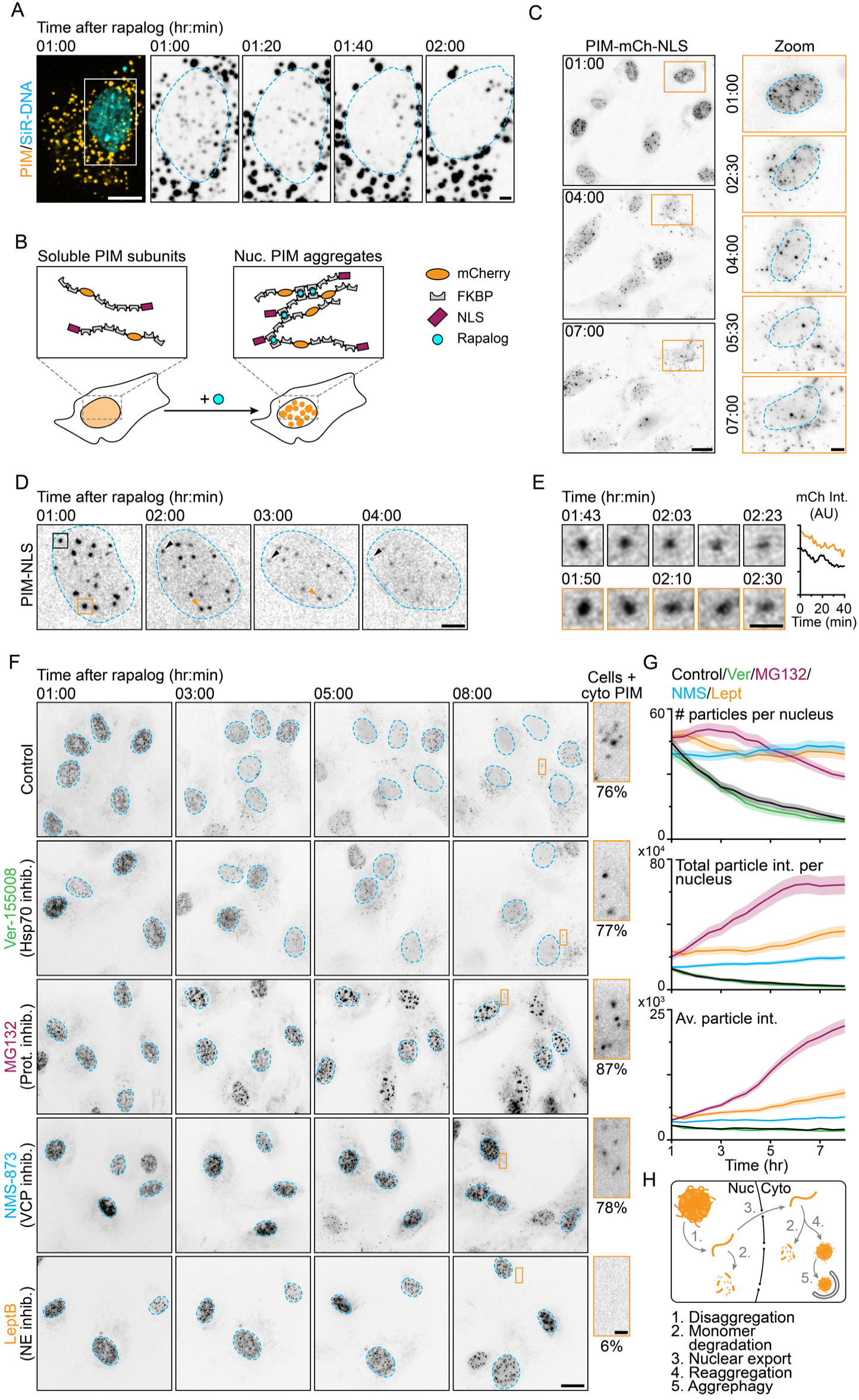
Nuclear PIM-mCh-NLS aggregates are cleared via a proteasome/VCP dependent mechanism. (**A**) Representative images of live-cell confocal imaging of U2OS-WT cells transfected with dual-PIM (orange), showing removal of nuclear PIM aggregates. The nucleus is visualized using SiR-DNA (cyan). (**B**) Schematic representation showing nuclear PIM as a tool to monitor nuclear disaggregation. Expression of PIM-mCh-NLS followed by rapalog addition yields nuclear PIM aggregates that can be monitored using fluorescence microscopy. (**C**) Representative images of RPE-PIM-mCh-NLS cells showing the gradual removal of nuclear of PIM-mCh-NLS aggregates (inverted contrast grey scale) and concomitant appearance of cytosolic aggregates. Zooms outlined in orange show nuclear PIM removal in a single RPE-PIM-mCh-NLS cell. (**D**) Sum projections of z-stacks acquired using live-cell confocal microscopy of RPE-PIM-mCh-NLS cells. (**E**) Zooms of D showing gradual decline of mCherry signal of single aggregates. (**F**) Representative images of RPE-PIM-mCh-NLS treated with Ver-155008, MG132, NMS-873 or LeptB. Orange boxes indicate a cytosolic region that is shown as a zoom, highlighting presence or absence of cytosolic aggregates after 8 hours. The percentages represent the fraction of cells with cytosolic aggregates. (**G**) Quantification of (**H**) showing the number of particles per nucleus (top graph), total intensity of particles per nucleus (middle graph) or the average particle intensity per nucleus (bottom graph) over time. Traces represent the mean and shaded areas show the SEM of control (grey), Ver-155008 (green), MG132 (magenta), NMS-873 (cyan) and LeptB (orange; N≥3, n_cells_ = 58, 74, 63, 66, respectively). Nuclei are outlined in cyan (C, D and F). Scale bars represent 20 μm (C and F), 5 μm (A, C_zoom_ and D) and 2 μm (E and F_zoom_).

Doxycycline induced expression of PIM-mCh-NLS resulted in homogeneous expression levels with minimal observable pre-aggregation (Fig. S1, A). As expected, addition of rapalog induced formation of nuclear aggregates within 1 hour (Fig. 1, C). Nuclear PIMs formed in the nucleoplasm and were excluded from the nucleolus (Fig. S1, B). Subsequent long term live-cell imaging revealed the gradual clearance of aggregates from the nucleus (Fig. 1, C). Strikingly, removal of nuclear PIM aggregates from the nucleus correlated with the appearance of cytosolic mCherry clusters that occasionally colocalized with the autophagosomal marker LC-3, showing that these aggregates are targets of autophagy (Fig. 1, C and Fig. S1, C). To visualize the removal of single aggregates from the nucleus, we used confocal imaging and visualized individual aggregates over time. We found that single aggregates were gradually dissolved, as shown by a gradual decline of mCherry fluorescence of aggregates (Fig. 1, D and E). Together, these observations suggest that nuclear aggregates are disaggregated through resolubilizing of PIM-mCh-NLS subunits that are subsequently exported from the nucleus and reaggregate in the cytosol.

### Nuclear PIM-mCh-NLS aggregates are cleared via a proteasome/VCP dependent mechanism

We set out to further study these observations in order to (1) study the mechanisms driving removal of nuclear PIM aggregates by probing nuclear PQC, and (2) elucidate the process underlying cytosolic reaggregation. To probe the protein quality control factors involved in the clearance of nuclear PIMs, we employed long term live-cell imaging of the RPE-PIM-mCh-NLS cells in the presence of various inhibitors (Fig. 1, F). We then used automated detection of single aggregates in whole nuclei to analyse the number of aggregates and total aggregate intensity per nucleus, and the average intensity of all PIM-mCh-NLS cluster in single nuclei (Fig. 1, G). As expected, control cells showed gradual removal of PIM-mCh-NLS aggregates from the nucleus (Fig. 1, F and G). After 8 hours, 47.0 % of cells had no, or very few (<5) nuclear aggregates. To our surprise, treating cells with Ver-155008 to inhibit Hsp70, a chaperone widely recognized as a chief disaggregase in mammalian cells, did not inhibit the clearance of PIM-mCh-NLS aggregates.

We therefore tested whether inhibition of alternative PQC proteins might affect disaggregation of nuclear PIMs. In contrast to inhibition of Hsp70, inhibiting the proteasome (using MG132) completely abrogated nuclear PIM clearance (Fig. 1, F and G). In this condition, nuclear PIMs remained present during imaging (8 hours) and appeared to cluster, since analysis revealed a modest reduction in the number of PIMs, but a strong increase in the total nuclear intensity and the average intensity of remaining particles (Fig. 1, F and G). Interestingly, addition of NMS-873 (VCP inhibitor) also prevented nuclear PIM clearance. Cells that were treated with VCP inhibitor showed persistent numbers of aggregates, even after 8 hours (Fig. 1, F and G). Lastly, we treated cells with leptomycin B (LeptB), a selective inhibitor of CRM1-dependent nuclear export. Inhibition of nuclear export also resulted in reduced nuclear aggregate clearance with aggregates being phenotypically similar to those found in VCP treated cells.

In addition to the striking removal of aggregates from the nucleus, the appearance of cytosolic PIMs was also impacted by drug treatment. Control cells, and cell treated with Hsp70, VCP or proteasome inhibitors, all showed accumulation of cytosolic aggregates 8 hours after rapalog treatment (78, 77 and 87% of cells with cytosolic aggregates, respectively, Fig. 1, F). In contrast, inhibiting nuclear export drastically reduced the number of cells with cytosolic aggregates (6%, Fig. 1, F), suggesting cytosolic aggregation indeed depends on PIM subunits originating from the nucleus.

Taken together, these results implicate VCP and the proteasome as key players in nuclear disaggregation and clearance of PIM aggregates. Furthermore, the reaccumulation of PIM aggregates in the cytosol suggests a mechanism of nuclear aggregate removal that involves nuclear disaggregation and nuclear export, followed by cytosolic reaggregation and subsequent autophagic clearance (Fig. 1, H).

### Targeted depletion of PQC subunits shows VCP, the 20S and 19S proteasome and DNAJB1 are involved nuclear disaggregation

While chemical inhibition of multi-subunit protein complexes or PQC networks, such as the proteasome or the Hsp70 triad, provides valuable insight into aggregate clearance mechanisms, it does not resolve contributions of individual protein members. Therefore, we employed siRNA knockdown of the various PQC protein subunits implicated in nuclear PIM clearance (Fig. 1, F).

Using siRNAs, we targeted the proteasome complex by knocking down 20S CP subunit PSMB2 and the 19S subunits PSMC2 and PSMC5 (Fig. S3). Additionally, we probed the involvement of Hsp70 and two J-domain proteins (DNAJB1 and DNAJB6) with broad functions in substrate recognition and disaggregation (Gao et al., 2015; Hageman et al., 2010; Kampinga & Craig, 2010; McMahon et al., 2021; Nillegoda et al., 2015; Serlidaki et al., 2020). To validate our findings that show chemical inhibition of VCP reduces PIM clearance, we also targeted VCP (Fig. 2, A). We performed knockdown of these targets in RPE-PIM-mCh-NLS cells, performed disaggregation assays, and fixed cells 3.5 or 6 hours after of rapalog addition (Fig. 2, B and C). Subsequent analysis of the aggregation coefficient, defined as the standard deviation of pixel intensity normalized by the mean pixel intensity, of individual nuclei was then used to visualize nuclear PIM aggregation levels in various conditions (Fig. 2, D).

**Figure 2:**
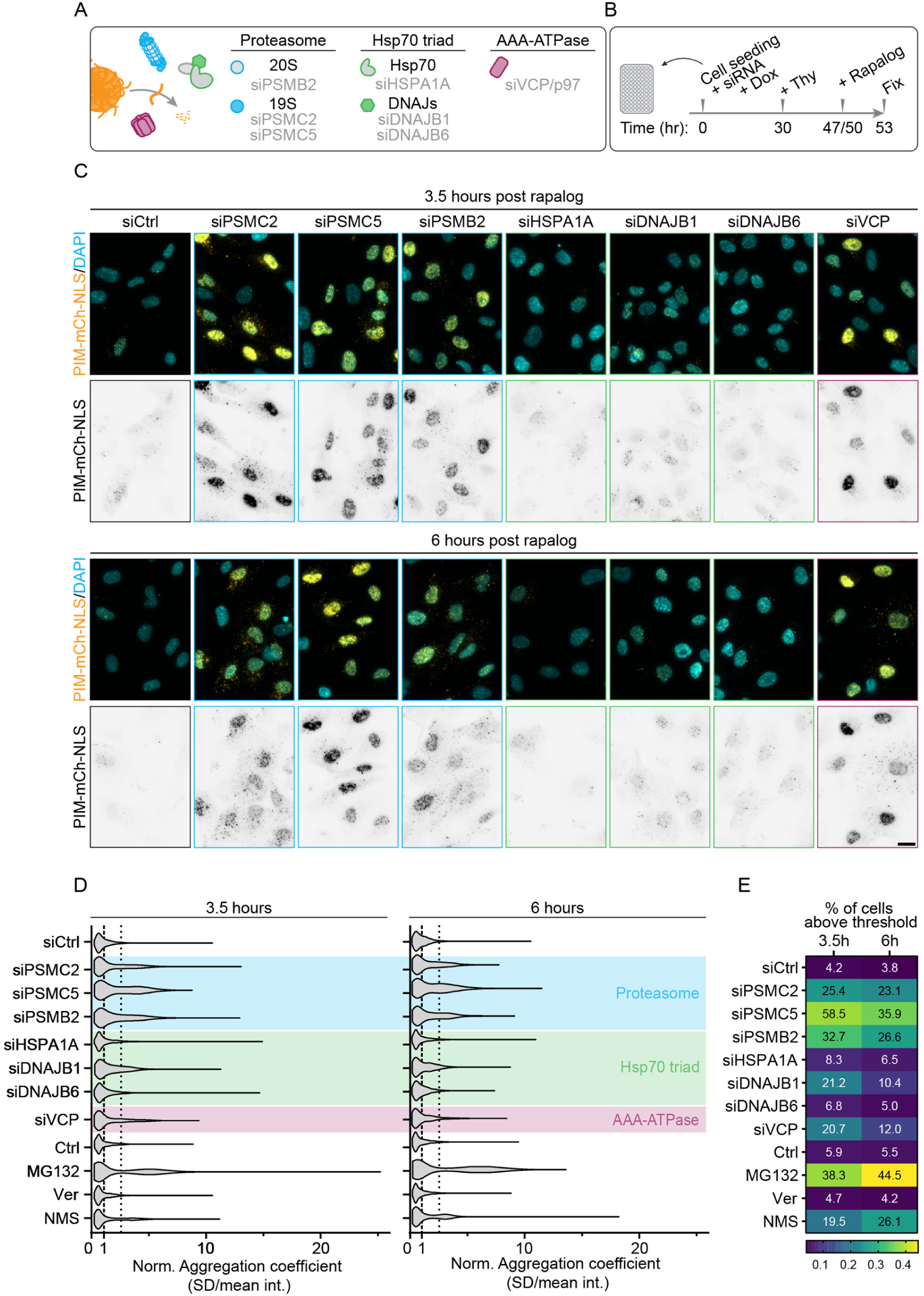
Targeted depletion of PQC players shows VCP, the 20S and 19S proteasome and DNAJB1 are involved nuclear disaggregation. (**A**) Schematic representation of siRNAs to target proteasome subunits (PSMB2, PSMC2 and PSMC5; cyan), Hsp70 triad members (HSPA1A, DNAJB1 and DNAJB6; green) and the AAA-ATPase VCP (siVCP/p97; magenta). (**B**) Schematic depicting the experimental setup prior to fixation, including induction of PIM-mCh-NLS expression, siRNA knockdown, cell cycle arrest (+Thy) and rapalog treatment. (**C**) Representative images of fixed RPE-PIM-mCh-NLS cells subjected to siRNA knockdown conditions previously summarized in (A). Cells were fixed 3.5 (top panel), or 6 (bottom panel) hours after aggregate induction using rapalog. (**D**) Quantification of the aggregation coefficient (SD/mean intensity per nucleus) of cells shown in (C) treated with si-Control, si-PSMC2, si-PSMC5, si-PSMB2, si-HSPA1A, si-DNAJB1, si-DNAJB6 or si-VCP (N=2, n_cells-3h_ = 846, 950, 926, 1012, 1339, 981, 1151, 703, respectively; n_cells-6h_ = 596, 901, 763, 864, 1175, 1078, 955, 315). Vertical lines represent normalized mean of siCtrl (1; dark grey) and 2 times SD. Shaded areas represent various target groups. Micrographs of non-siRNA treatment conditions are shown in fig. S2. (**E**) Graph depicting the percentage of cells with an aggregation coefficient above the cutoff value. Scale bar represent 20 μm.

As expected, nuclear PIMs in cells treated with non-targeted siRNA were mostly cleared, with some cells harbouring low amounts of small aggregates 3.5 or 6 hours after onset of aggregation. Interestingly, knock down of either 20S or 19S proteasomal subunits resulted in high aggregation levels of PIMs, comparable to those found in cells treated with MG132 (Fig. 2, C and D, and Fig. S2). Knock down of VCP also strongly inhibited disaggregation and removal of nuclear PIMs, similar to addition of NMS-873. In contrast, targeting Hsp70 (siHSPA1A) did not reduce PIM clearance, and treatment with Hsp70 inhibitor showed a similar phenotype (Fig. 2, C and D, and Fig. S2), validating previous live-cell imaging experiments (Fig. 1, F and G). Interestingly, cells subjected to DNAJB1 knock down showed low levels of nuclear PIM aggregation at both time points, but these levels appeared slightly elevated compared to control siRNA. In contrast, targeting of DNAJB6 had no effect on PIM clearance. Quantification of the fraction of cells with a high aggregation coefficient compared to control siRNA again showed a potent role for the proteasome, with knock down of 19S subunit PSMC5 most strongly affecting PIM aggregation at both timepoints, and showed variation in the contribution of J-domain proteins (Fig. 2, E).

Thus, knock down of proteasomal subunits and VCP strongly inhibits nuclear PIM clearance, validating our earlier findings using inhibitors. In addition, the 19S and 20S proteasome subunits are both required for efficient disaggregation of PIMs. Furthermore, these results also suggest DNAJB1 is involved in PIM disaggregation or in the prevention of PIM aggregation in an Hsp70 independent manner.

Having identified VCP as a major player in nuclear clearance of PIMs, we asked whether nuclear clearance could be enhanced via the overexpression of either VCP, a disease-associated VCP mutant (VCP(R155H)) or different cofactors (JAB1, UBXD1, VCF1). Whereas in most conditions there was little impact on nuclear PIM clearance (Fig. S4), overexpression of VCF1 (VCP Nuclear Cofactor Family Member 1) strongly inhibited nuclear clearance (Fig. S4). VCF1 promotes VCP-UFD1-NPL4 interactions with substrates and has an unusually high binding affinity for VCP (Mirsanaye et al., 2024). This suggests that nuclear disaggregation is independent of VCP’s interaction with UFD1 and NPL4 and that VCF1 overexpression outcompetes other VCP cofactors needed for disaggregation.

### PIM-mCh-NLS monomers reaggregate in the cytosol after nuclear disaggregation

The redistribution of PIMs to the cytosol is a striking phenotype consistently observed in our assays (Fig. 1, C and F). Interestingly, exchange of client molecules to PQC machinery in different cellular compartments has been observed in various studies. Specifically, targeting of cytosolic substates for proteasomal degradation in the nucleus is well studied in yeast (Banik et al., 2024; Jones & Gardner, 2016; Prasad et al., 2010, 2018; Shakya et al., 2021), and is likely conserved in mammalian cells (S. H. Park et al., 2013; S. J. Park et al., 2024). Conversely, some nuclear proteins are targeted to the cytosol for degradation (Dou et al., 2015). However, to our knowledge, nuclear dissolution and subsequent cytosolic reaggregation of aggregation prone proteins following nuclear export has not been reported previously.

To further study the redistribution of PIM aggregates, we aimed to determine whether the cytosolic aggregates formed in our cells indeed consist of subunits originating from nuclear aggregates. While the observed lack of cytosolic redistribution in LeptB treated cells strongly suggests the nuclear origin of cytosolic PIMs in our assay, we wondered whether the formation of cytosolic aggregates after rapalog addition could also be the result of the aggregation of newly synthesized PIM-mCh-NLS monomers. In order to test this hypothesis, we blocked *de novo* protein synthesis, either by washing out doxycycline for 24 hours or by treating cells with cycloheximide (CHX, Fig. 3, A). We found that PIM aggregates still emerged in the cytosol under these conditions, indicating that cytosolic PIMs are formed by pre-existing PIM monomers.

**Figure 3:**
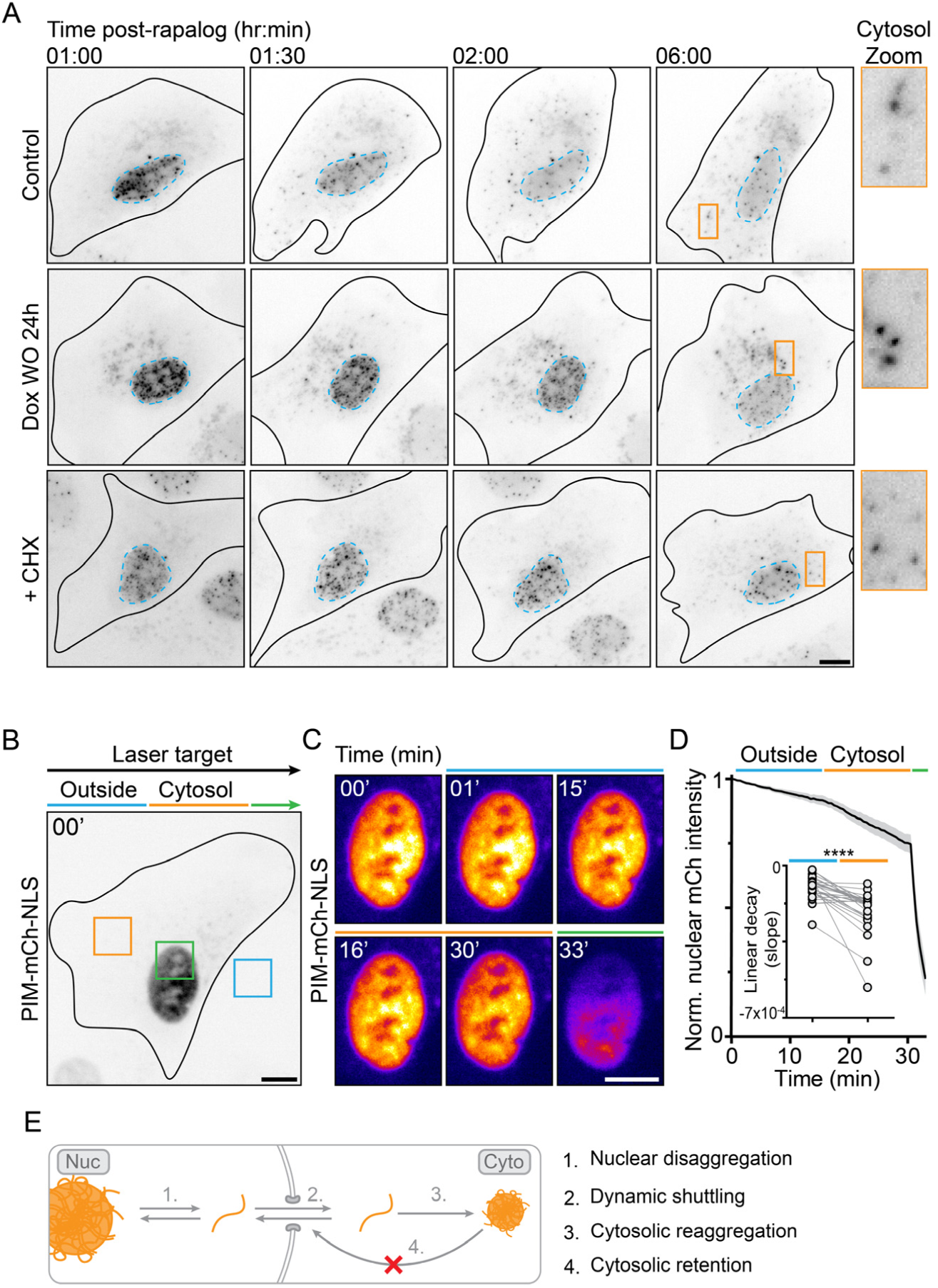
PIM-mCh-NLS monomers reaggregate in the cytosol after nuclear disaggregation. (**A**) Representative live-cell microscopy images of control RPE-PIM-mCh-NLS cells, cells subjected to doxycycline washout (Dox WO) and cells treated with cycloheximide (CHX). Zooms (orange boxes) indicate presence of cytosolic PIM aggregates after 5 hours. Nuclei are outlined in cyan. (**B**) Representative z-projection of RPE-PIM-mCh-NLS cell imaged without rapalog addition. Arrows indicate photobleaching strategy sequentially targeting a extracellular (cyan), cytosolic (orange) or nuclear (green) ROI. (**C**) Zoom of the nucleus of the cell shown in (B) showing soluble PIM-mCh-NLS intensity during various photobleaching regimes. (**D**) Quantification of the normalized nuclear mCherry intensity of cells shown in (C; N=3, n=8, 10, 9). Trace represents the mean ± 95% confidence interval. Inset graph shows rate of decay of nuclear mCherry signal. Dots represent single cells. Connecting lines link repeated measurements and differences in decay rates were statistically tested using a Wilcoxon matched-pairs signed rank test. (**E**) Proposed model for nuclear removal of nuclear PIM aggregates, that requires (1) nuclear disaggregation, (2) dynamic nucleocytoplasmic shuttling, (3) cytosolic reaggregation, and (4) retention by size exclusion. **** = p < 0.0001. All scale bars represent 10 μm.

How then are these solubilized PIM monomers translocated to the cytosol? To answer this question, we studied the nucleocytoplasmic shuttling dynamics of non-aggregated PIM subunits. To this end, we imaged cells expressing PIM-mCh-NLS without adding rapalog and sequentially photobleached an extracellular and cytosolic ROI. To determine the efficacy of our photobleaching, we ended each acquisition by bleaching a ROI placed in the nucleus (Fig. 3, B). This approach enabled us to determine whether soluble PIM-mCh-NLS undergoes nucleoplasm shuttling or remains strictly nuclear, because dynamic exchange of PIM-mCh-NLS between the nucleus and cytosol would enable us to bleach the nuclear mCherry signal by targeting a cytosolic ROI. We found that PIM-mCh-NLS subunits are indeed dynamically exchanged between the nucleus and the cytosol, since photobleaching of the cytosol resulted in an increased decay rate of nuclear mCherry signal compared to extracellular photobleaching at a similar distance to the nucleus (Fig. 3, C and D). Together with our previous findings, these data indicate that cytosolic reaggregation depends on nuclear export and not on *de novo* protein synthesis, and suggest that cytosolic aggregates form after solubilized PIM monomers exit the nucleus via dynamic nucleocytoplasmic shuttling. Subsequent cytosolic reaggregation would then ensure cytosolic retention of PIM-monomers, followed by autophagy (Fig. 3, E).

### Structural modelling reveals accessible NES in unfolded PIM-mCh-NLS

Given that the PIM subunits appearing in the cytosol are of nuclear origin, what triggers their export? The inhibition of this redistribution in LeptB-treated cells suggests a strong preference for nuclear export of newly disaggregated subunits and implicates the shuttling factor CRM1 as the driving factor. We therefore wondered whether the potential unfolding of PIM-mCh-NLS during disaggregation might alter its shuttling dynamics.

To explore PIM-mCh-NLS interactions with shuttling factors, we used NLSExplorer (Y. F. Li et al., 2025), NLStradamus (Nguyen Ba et al., 2009), and NESmapper (Kosugi et al., 2014) to predict possible nuclear import and export signals within the PIM-mCh-NLS sequence. As expected, NLS predictions revealed only the known sequence at the end of the construct (Table S1). Interestingly, nuclear export signal (NES) predictions revealed 12 NES candidates: two per FKBP homodimerization domain, dubbed NES1 and NES2 (the second NES within the second FKBP region contains a single amino acid difference and so is regarded as NES2B; Fig. 4, A, Table S2).

**Figure 4:**
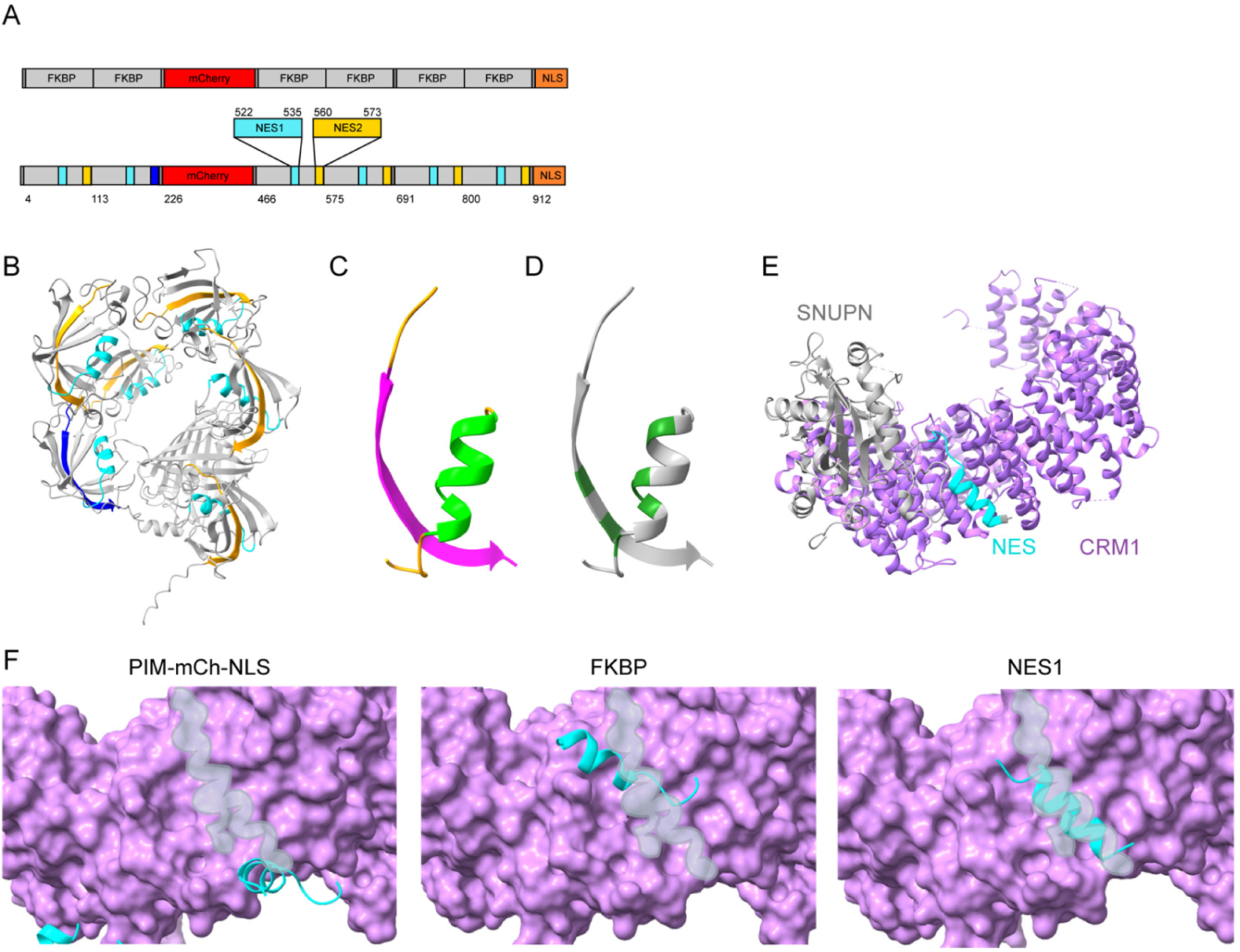
Structural modelling suggests NES unshielding upon PIM-mCh-NLS unfolding. (**A**) Schematic representations showing PIM-mCh-NLS protein domains and NESmapper-predicted NES motifs. (**B**) AlphaFold3-predicted structure for PIM-mCh-NLS coloured to show NES1 (cyan), NES2 (yellow), NES2B (blue). (**C**) Zoom showing predicted NESs in single FKBP homodimerization domain. Coloured to show secondary structure (NES1: β-strand, magenta; NES2: α-helix, light green; coil, orange). (**D**) Re-coloured to show buried surface area (dark green) as calculated by ChimeraX. (**E**) Crystal structure of CRM1-SNUPN complex (CRM1, purple; SNUPN, grey; SNUPN NES motif, cyan) obtained from UniProt. (**F**) Protein-protein docking for CMR1 (purple) and PIM-mCh-NLS, isolated FKBP domain, or isolated NES (NES1 regions in cyan) predicted by HADDOCK. CRM1’s hydrophobic binding pocket overlaid in grey.

Because sequence-based predictions are prone to false-positives (Y. Lee et al., 2019), we performed further structural validation of the NES candidates. Most NES motifs adopt an α-helix (or α-helix extended) conformation to correctly dock into the hydrophobic groove of CRM1 (Dong et al., 2009; Fung et al., 2015, 2017). We used AlphaFold 3 (Abramson et al., 2024) to predict the folded structure of PIM-mCh-NLS (Fig. 4, B). NES1 formed an α-helix, suggesting it could theoretically bind to CRM1 (Fig. 4, C). NES1 further maintained the α-helix structure when modelled in isolation (Fig. S5, A), suggesting it could locally adopt this structure when the monomer was unfolded (such as during disaggregation). NES2 (and NES2B) formed part of a β-sheet and did not fold when modelled in isolation (Fig. S5, B), making it highly unlikely to bind to CRM1 (Fig. 4, C).

An NES motifs’ accessibility for CRM1 binding depends upon its position within a protein: ideally an NES should be located at either the N- or C-terminus, or within an unstructured region (Y. Lee et al., 2019). NES1 is located within the folded FKBP domain, making it a poor contender for CRM1 binding. This was supported by calculating the buried and solvent-accessible surface areas using ChimeraX (Meng et al., 2023), which confirmed that NES1 was partially buried (Fig. 4, D). These results suggest that, within a normally folded PIM monomer, NES1 cannot bind CRM1, but that unfolding the monomer (such as by VCP) would reveal the buried NES.

To confirm that NES1 could only bind to CRM1 when the PIM monomers are unfolded, we used the HADDOCK (High Ambiguity Driven protein-protein DOCKing) server (Honorato et al., 2021, 2024) to predict how either full-length PIM-mCh-NLS, a single FKBP domain, or an NES1 in isolation would dock to free CRM1 (Berman et al., 2000; Monecke et al., 2013). These predictions were compared against the crystal structure of CRM1 bound to SNUPN as a control (Berman et al., 2000; Monecke et al., 2013) (Fig. 4, E). When simulating full-length PIM-mCh-NLS or a single FKBP domain, no conformation was achieved that approximated the canonical CRM1 binding, but when NES1 was modelled in isolation it was able to dock into the hydrophobic groove of CRM1 (Fig. 4, F). Together, these results support a model in which the PIM monomers contain a buried NES that is revealed through protein unfolding, which then drives nuclear export after disaggregation.

### Monomer resolubilization is prioritized over proteolysis

The finding that disaggregated PIM subunits from the nucleus can reaggregate in the cytosol, suggests that the clearance of PIMs mostly relies on disaggregation and export, rather than proteolysis. These findings appear in contrast with our earlier findings in which nuclear PIM clearance is efficiently inhibited by targeting catalytic activity of the proteasome. However, the level of degradative capacity of nuclear proteasomes is debated (Dang et al., 2016; Enam et al., 2018), and recent *in vitro* evidence suggests the proteasome can fragment fibrillar protein assemblies independent of its proteolytic activity (Cliffe et al., 2019). Moreover, the 19S subunit contributes to fragmentation of cytosolic aggregates prior to autophagic clearance (Mauthe et al., 2025). Therefore, we set out to further study the contribution of protein degradation during disaggregation in the nucleus.

To this end, we realized that treatment with LeptB can be used to exclusively study nuclear processing of PIM-mCh-NLS by eliminating nuclear export and cytosolic processing. First, we tested whether disaggregated PIM subunits could reaggregate by expressing a variant of the PIM construct containing photoactivatable GFP (PAGFP, PIM-mCh-PAGFP-NLS, Fig. 5, A) in U2OS WT cells. We treated these cells with rapalog and LeptB, to maintain stable amounts of nuclear aggregates without abolishing PIM disaggregation. As expected, rapalog addition resulted in the formation of nuclear aggregates that were mCherry positive but did not emit PAGFP signal upon blue light illumination (Fig. 5, B). Using sequential activation of equally sized areas containing either nucleoplasm or PIM aggregates, we found that aggregated PIM-mCh-PAGFP-NLS subunits can indeed be resolubilized and subsequently reaggregate into other PIM aggregates (Fig. 5, B and C). The contribution of activation of PIM-mCh-PAGFP-NLS subunits that were already in solution upon photoactivation is likely limited, since only activation of PIM aggregates, and not nucleoplasmic regions containing few solubilized PIM subunits resulted in a strong accumulation of PAGFP signal in non-activated PIM aggregates (Fig. 5, C). Together, these results indicate that PIM disaggregation is not tightly coupled to proteolytic degradation, but generates PIM species that can re-aggregate.

**Figure 5:**
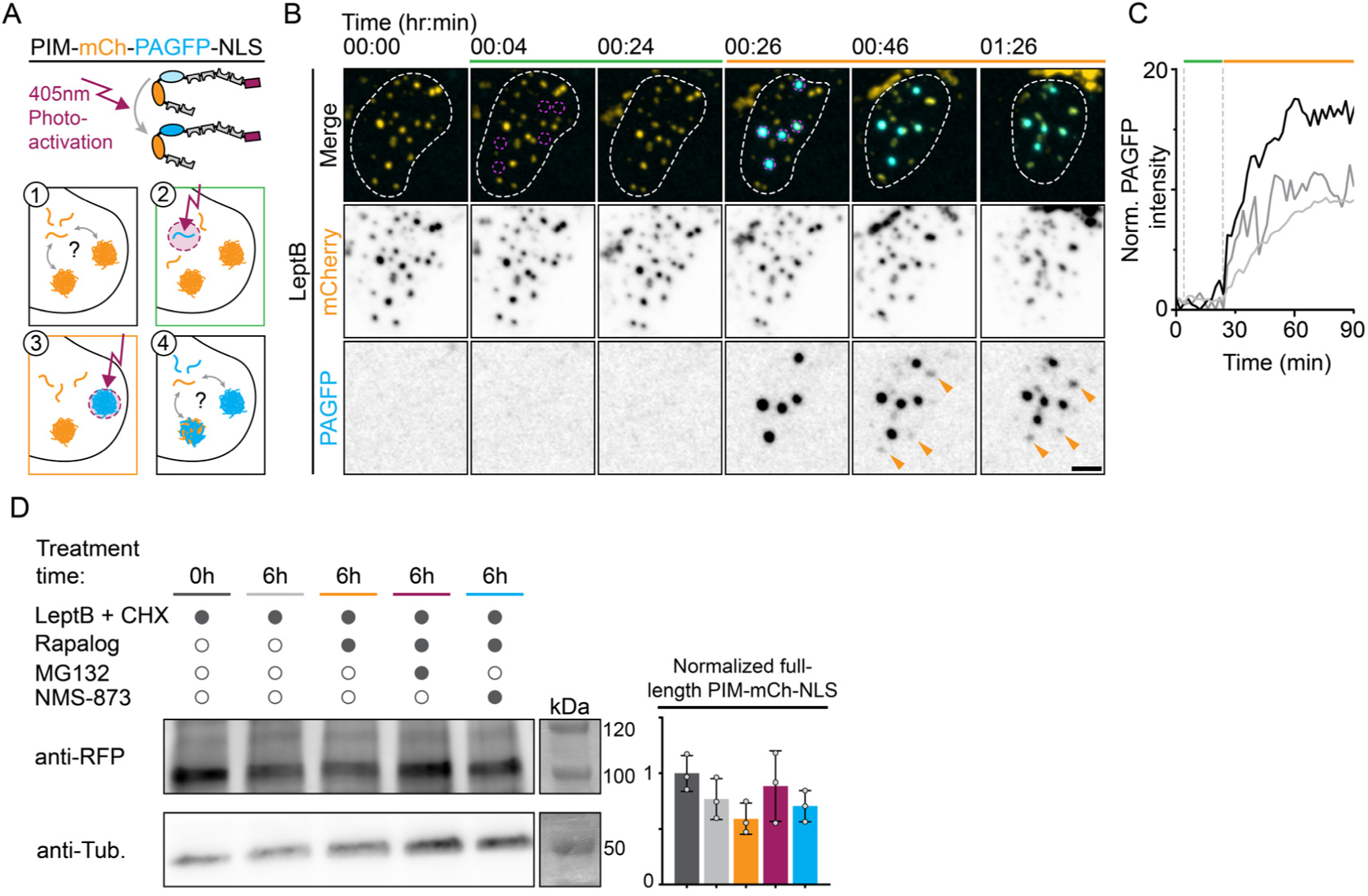
Monomer resolubilization is prioritized over proteolysis. (**A**) Graphic representation of experiment performed in U2OS-WT cells expressing PIMs harbouring photoactivatable GFP (PIM-mCh-PAGFP-NLS; 1). Sequential activation of PAGFP in the nucleosol (2) and in aggregates (3) and subsequent monitoring of PAGFP signal (4). (**B**) U2OS-WT cells expressing PIM-mCh-PAGFP-NLS and treated with LeptB for ∼ 2 hours. Cells were subjected to sequential photoactivation of PAGFP as shown in (A). Orange arrowheads indicate accumulation PAGFP in non-activated nuclear PIM aggregates and dashed line delineate the nucleus. (**C**) Graph showing normalized PAGFP intensity of non-activated nuclear PIM aggregates indicted by arrowheads in (B). (**D**) Representative images of western blot of the lysate of cells treated with nuclear export inhibitor (LeptB) and protein synthesis block (CHX), and rapalog, MG132 or NMS-873. Membranes were probed using anti-RFP and anti-α-Tubulin antibodies. Graph shows normalized intensity of full lengths PIM-mCh-NLS. Dots are independent replicates (N=3), bars represent the mean, and error bars indicate the SD. Scale bar represents 5 μm.

To further probe the extent to which nuclear proteasomal degradation affects PIM-mCh-NLS clearance we took a biochemical approach to determine the total levels of full-length PIM-mCh-NLS under various conditions. We again performed nuclear export block in all conditions to selectively study the nuclear processing of PIM subunits. Addition of CHX to block protein synthesis enabled us to monitor the amount of full-length PIM-mCh-NLS over time. In these conditions, RPE-PIM-mCh-NLS cells were subjected to control treatment (DMSO) for 0 or 6 hours, or treated with MG132 or NMS-873 for 6 hours. Protein detection using western blot revealed a baseline turnover of unaggregated PIM-mCh-NLS subunits of ∼23.1% during 6 hours (control 0 hours vs. control 6 hours; Fig. 5, D and Fig. S6). Compared to control, PIM degradation was only slightly increased in cells that were treated with rapalog (17.6% reduced compared to the 6 hour control). The increased PIM turnover observed upon rapalog addition could be rescued by the addition of proteasome inhibitor MG132 (26.4% higher compared to rapalog-treated cells), suggesting that proteasomal proteolysis is only partially responsible for the degradation of PIM monomers. Treating cells with the VCP inhibitor NMS-873 slightly increased the amount of signal observed after 6 hours, but the high variability between replicates precluded strong conclusions. Nonetheless, together these results indicate that many monomers are not degraded in the nucleus following PIM disaggregation.

### Proteasome inhibition limits the dynamic turnover of nuclear aggregates

Our results identify both VCP and the proteasome as crucial players in nuclear PIM clearance, and suggest a limited role of proteolysis during nuclear PIM clearance. However, it remains unclear how these two players would cooperate to facilitate efficient nuclear disaggregation. Previous work that identified disaggregase activity of VCP often links its function to subsequent proteasomal degradation of substrates (Nillegoda et al., 2018; Phan et al., 2024; Saha et al., 2023). However, our current findings, in combination with previous work showing proteasome independent fragmentation that does not rely on proteolysis (Cliffe et al., 2019), might suggest alternative (cooperative) mechanisms employed for nuclear disaggregation of PIMs.

To further study VCP- and proteasome-mediated removal of PIMs, we probed aggregate dynamics using high resolution, live-cell microscopy and FRAP (Fluorescence Recovery After Photobleaching). Using confocal microscopy, we visualized nuclear aggregates in RPE-PIM-mCh-NLS cells treated with LeptB, NMS-837, or MG132 for an extended time (∼ 4 hours). In line with previous imaging, we found that proteasome inhibition using MG132 yielded large aggregates. Interestingly, MG132 treatment also caused more amorphous aggregates that often appeared as clusters of multiple aggregates (Fig. 6, A). In contrast, cells treated with nuclear export block or VCP inhibitor showed smaller globular clusters. To determine the dynamics of PIM aggregates in these conditions, we performed FRAP on individual aggregates and monitored post-bleaching recovery for just under 90 minutes (Fig. 6, B). Aggregates in LeptB treated cells showed substantial recovery that eventually surpassed their original signal intensity, indicating ongoing disaggregation and reaggregation (127.6±47.5% recovery after 88.5; Fig. 6, C). Inhibiting the proteasome or VCP reduced this recovery to different degrees (51.6±26.7% recovery after 88.5 minutes for proteasome inhibition, 84.5±36.0% for VCP inhibition). Interestingly, the reduced recovery upon VCP inhibition was only apparent during later time points and the early fluorescence recovery closely matched that with LeptB (NMS 58.8±20.8% recovery after 30 minutes; LeptB 66.6±22.1% recovery after 30 minutes). We did not observe an increase in the intensity of non-bleached aggregates for the first 45 minutes of recovery in any of our experimental conditions, which suggests that ongoing dynamic exchange, and not net growth, is causing the recovery during this time period (Fig. 6, C). However, after 45 minutes the LeptB condition did exhibit an increase in intensity for non-bleached aggregates, and accounting for this increase eliminated the difference between the LeptB and NMS-873 conditions (Fig. 6, C). Thus, our findings reveal a role for the proteasome in aggregate turnover and also demonstrate continuing disaggregation upon VCP inhibition.

**Figure 6:**
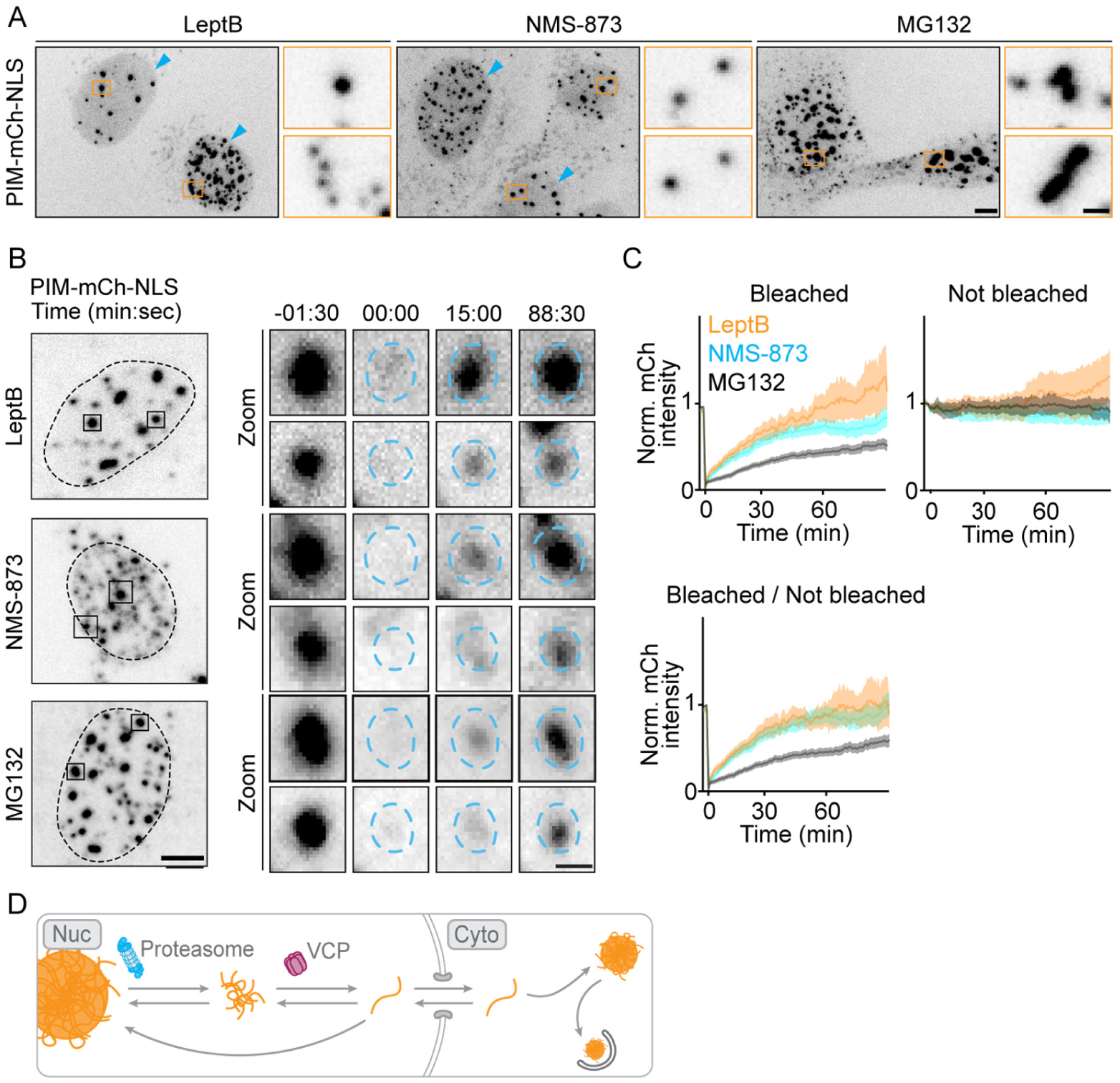
Proteasome inhibition limits the dynamic turnover of nuclear aggregates. (**A**) Representative images of RPE-PIM-mCh-NLS cells treated with rapalog for ∼4 hours, that were also treated with LeptB, NMS-873 or MG132. Zooms (orange boxes) show aggregate shape and size in various conditions. Cyan arrowheads indicate high nucleosolic mCherry signal. (**B**) Representative images of RPE-PIM-mCh-NLS cells treated as in (A). Dotted line indicates the nucleus. Zooms (black boxes) indicate representative aggregates targeted for FRAP. (**C**) Graph showing the average normalized mCherry intensity of aggregates after photobleaching (FRAP) in conditions shown in (B). Traces represent the mean, shaded areas show the 95% confidence interval (N>3, n_PIMs_=35_LeptB_, 43_NMS-873_, and 67_MG132_). (**D**) Proposed multistep mechanisms for nuclear aggregate clearance. Scale bars represent 5 μm or 1 μm (Zooms).

Closer examination of these conditions revealed a strong increase in nucleosolic mCherry in conditions that showed high levels of FRAP recovery (LeptB and NMS-treated cells, Fig. 6, A). Such an increased level of nucleosolic signal was expected in LeptB treated cells, because it indicates ongoing disaggregation without nuclear export. A similar increase of the nucleosolic signal upon VCP inhibition is more surprising and also indicates an impaired export of PIM-mCh-NLS following disaggregation. These observations are consistent with a multistep model in which VCP acts downstream of the proteasome and disaggregates small PIM fragments generated by proteasomal fragmentation of the original PIMs (Fig. 6, D). Such fragments would be too small and mobile to be resolved by conventional light microscopy techniques but would still exceed the size restriction of the nuclear pore, which explains the increased nucleosolic mCherry signal.

## Discussion

Here we generated a robust cellular assay that enables induction of nuclear protein aggregation and employed it to study the mechanisms of nuclear aggregate removal. We found that nuclear PIM aggregates are rapidly cleared from the nucleus (∼5 hours) by disaggregation and nuclear export of soluble monomers. The removal of PIMs depended on VCP and the proteasome but was strikingly unaffected by inhibition or knock down of Hsp70. Subsequent CRM1 dependent nuclear export caused reaggregation of nuclear PIM subunits in the cytosol. Our work thus highlights the differential handling of identical substrates by nuclear and cytosolic PQC machinery. Whereas cytosolic aggregates are slowly degraded through autophagy, nuclear aggregates are rapidly resolubilized, resulting in cytosolic reaggregation and subsequent clearance through autophagy. It thus appears that the inability to perform autophagy in the nucleus has promoted the development of a highly effective nuclear disaggregation machinery. Targeting the proteasome, via knockdown or using chemical inhibition, consistently resulted in a strong decrease in PIM clearance. Interestingly, the contribution of proteolysis appears to be limited in our assays. Therefore, our data strongly suggest that the proteasome functions as an alternative disaggregase. This is in line with *in vitro* work by Cliffe *et al*. that shows the 26S proteasome can fragment filamentous aggregates, such as tau fibrils (Cliffe et al., 2019). Importantly, this fragmentase function depends on the complete holoenzyme (20S and 19S) and the 19S ATPase activity, but was shown to be independent of catalytic activity of the 20S CP. Similarly, our si-RNA assay showed that nuclear PIM clearance depends on both the 20S CP, and the 19S subunit that has AAA-ATPase activity. However, we also find that PIM clearance is strongly affected by blocking the proteolytic activity of the proteasome (using MG132). This might suggest that the level of proteolysis depends on the aggregate substrate, but could also reflect a more general stalling of 26S proteasomes or PQC following MG132 treatment, potentially sequestering 19S subunits, and resulting in a loss of effective fragmentase function in cells. Fragmentation activity of the 19S proteasome was recently shown to be required for effective cytosolic clearance via aggrephagy in cells (Mauthe et al., 2025). Interestingly, this cytosolic fragmentation also requires Hsp70 and DNAJB6, while our assays identify VCP and DNAJB1 as potential contributors in nuclear fragmentation. Together, these findings show that 19S dependent fragmentation is a conserved pathway, with different co-regulators in the nucleus and cytosol.

We furthermore identified the involvement of VCP in the clearance of nuclear PIMs. Previous work already demonstrated that VCP can target cytosolic aggregates formed by tau, TDP-43, and Huntingtin (Benn et al., 2024; Darwich et al., 2020; Ghosh et al., 2018; Saha et al., 2023; Wani et al., 2021), but its role in nuclear disaggregation has remained unclear. Importantly, our work now shows that VCP contributes to clearance of nuclear aggregates, which is in line with another recent report showing VCP activation reduces levels of artificially induced nuclear TDP-43 aggregates (Phan et al., 2024). Interestingly, VCP mediated disaggregation is often linked to subsequent proteolytic breakdown of substrates (Gallagher et al., 2014; Meyer et al., 2012; Nillegoda et al., 2018; Saha et al., 2023). In contrast, the abundant redistribution of disaggregated subunits in our assay suggests limited proteolysis during clearance of nuclear PIMs. In addition, our FRAP analysis indicates that VCP most likely functions downstream of the proteosome and disaggregates small fragments generated by the proteosome to facilitate their export to the cytoplasm.

Importantly, while the assays developed in this work enable robust quantification of nuclear aggregate clearance, our light microscopy approach is limited in spatial and temporal resolution. In order to precisely determine the role of the proteasome and VCP in aggregate fragmentation and disaggregation, it would be highly beneficial to develop optimized biochemical and microscopy assays that enable detection of aggregate fragments that are currently unresolved. In addition, while our predictive analyses indicate the presence of a buried NES that is revealed during disaggregation, we did not validate this site through binding assays or mutational analysis. Further experiments could confirm the functionality of the predicted NES, and whether disrupting this site would inhibit cytoplasmic re-aggregation of PIMs.

Finally, we found that reducing Hsp70 activity has no effect on nuclear PIM clearance. However, our siRNA screen did reveal the Hsp70 co-chaperone DNAJB1 might have a modest role in PIM disaggregation. This finding suggest that substrate specificity of J-domain proteins, a well-recognized phenomenon for DNAJs in disaggregation and suppression of aggregation (Ayala Mariscal et al., 2022; Gao et al., 2015; Hageman et al., 2010; Kampinga et al., 2019; McMahon et al., 2021; Serlidaki et al., 2020), potentially extends to PIM aggregates. Since Ver-155008 also inhibits other members of the HSP70 family, such as Hsc70 (Schlecht et al., 2013), our data also suggest that J-domain protein action is HSP70-independent. Such a function is debated, but substrate binding by J-domain proteins can occur independently, which would enable HSP70-independent action (Kampinga et al., 2019; Kampinga & Craig, 2010).

In summary, our work establishes nuclear PIMs as a robust and versatile tool that enables high throughput analysis of PQC of nuclear aggregates. This revealed a novel route for nuclear aggregate removal, relying on disaggregation and nuclear export, followed by cytosolic reaggregation and clearance through autophagy. Together, this mechanism enables the clearance of nuclear substrates by autophagy.

## Acknowledgements

We thank Harm H. Kampinga and Mario Mauthe for fruitful discussions, and for the guidance on nuclear disaggregation mechanisms. We also thank Harm H. Kampinga for contributing reagents. We thank Sam van Beuningen for advice about high throughput screening approaches. Author contributions: G.K. and L.C.K. designed the research. G.K. created reagents, and G.K. and G.P.S. performed experiments, analysed data, and generated figures. A.F.J.J. designed and generated the PIM-mCh-NLS construct and performed pilot experiments. W.N. assisted in generation of stable cell lines and performed fixed-cell imaging of si-RNA experiments. G.K., G.P.S., and L.C.K wrote the manuscript with input from other authors. L.C.K. supervised the study.

## Material and Methods

### Constructs

Initial observations of rapid removal of nuclear aggregates were made using the pβactin-dualPIM construct containing several FKBP homo- and heterodimerization domains containing sequence variations, and mCherry and EGFP fluorophores (4x-FKBPhomo-cCherry-EGFP-2xFKBPhet; Addgene plasmid # 111758) (Janssen et al., 2018). To obtain nuclear targeting of this construct we added the NLS of the Sv40 large T-antigen (PKKKRKV) at the C-terminus, generating dualPIM-NLS. The construct used for photoactivation experiments (PIM-mCh-PAGFP-NLS) was created by substitution of EGFP for PAGFP.

To generate constructs for production of stable lines expressing PIM-NLS, we first removed the EGFP from the dualPIM-NLS construct. This vector was subsequently ligated into a pcDNA5-FRT-T0-empty-Puro vector after digestion with BamHI and HindIII (ER0051 and ER0505, respectively; Thermo Fisher), generating pcDNA-FRT-T0-PIM-mCherry-NLS. The pcDNA5-FRT-T0-empty-Puro vector was a kind gift from Geert Kops (Hubrecht Institute, Utrecht, the Netherlands).

### Generation of RPE-1-PIM-mCh-NLS line

For generation of RPE-1-Flp-In cells stably expressing PIM-mCherry-NLS, RPE-1-Flp-In-empty cells (a gift from Geert Kops) were co-transfected with pcDNA-FRT-T0-PIM-mCherry-NLS and pOG44 (Invitrogen) plasmids (3 μg DNA per well, ratio 1:9) using FuGENE6 (Promega) at a ratio of 3 μl reagent per 1 μg DNA. 2 days after transfection, the medium was supplemented with 0.0025 mg/ml puromycin.

### Cell lines and Cell culture

RPE-1-Flp-In-PIM-mCherry-NLS cells were cultured in DMEM/F12 (Gibco) with 10% FBS supplemented with 1% penicillin and streptomycin, and blasticidin (0,005 mg/ml) and puromycin (0,0025 mg/ml). U2OS WT cells were purchased from ATCC and were cultured in DMEM (Capricorn) supplemented with penicillin and streptomycin. Cells were maintained at 37°C and 5% CO2. Cells were regularly tested for mycoplasm.

### Transfection/Induction of PIM-NLS expression

For live cell experiments using U2OS WT cells, cells were seeded on 18 mm coverslips 1 day prior to transfection. Cells were subsequently transfected with 2 μg pβactin-dualPIM-NLS or pβactin-PIM-mCh-PAGFP-NLS using FuGENE6 (Promega) at a ratio of 3 μl reagent per 1 μg DNA. Cells were imaged ∼24 hours after transfection. In RPE-PIM-mCh-NLS cells PIM-NLS expression was induced by addition of 1 μg/ml doxycycline to the growth medium 1 day prior to the start of imaging. Just before the start of experiments, doxycycline was removed from the medium.

### Transfection of VCP and VCP cofactors

For fixed cell experiments, RPE-PIM-mCh-NLS cells were seeded on 18 mm coverslips 1 day prior to transfection. Cells were transfected with 1 μg of DNA using FuGENE6 (Promega6) at a ratio of 3 μl reagent per 1 μg DNA. For live cell experiments, cells were seeded on 25 mm coverslips and transfected with 2 μg of DNA. Cells were respectively fixed or imaged ∼24 hours after transfection. The following constructs were used: VCP(WT)-EGFP (Plasmid #23971, Addgene), VCP(R155H)-EGFP (Plasmid #23974, Addgene), UBXD1-GFP (Plasmid #86464, Addgene), EYFP-JAB1 (Plasmid # 111213), GFP-VCF1 WT (Plasmid #223018, Addgene).

### PIM formation and drug treatment

The formation of PIM aggregates was initiated by addition of 500 nM rapalog2 (635059, Takara) for 45 minutes and was subsequently washed out of the medium. Cells were then transferred to the microscope or incubated for live- or fixed- cell imaging at later timepoints. For all experiments that involved drug treatment, drugs were added 30 minutes before addition of rapalog2 and were replenished after rapalog2 washout. Drugs were added at the following concentrations; 10 μM Ver-155008 (SML0271, Sigma), 10 μM MG132 (474790, Calbiochem), 2 μM NMS-873 (SML1128, Sigma), 20 nM leptomycin B (LKT-L1761-C005, LKT). For high resolution imaging of cells expressing dualPIM, cells were incubated with 200 nM SiR-DNA (SC015, Spirochrome) as a nuclear marker.

For western blot analysis of nuclear PIM-mCh-NLS breakdown (Fig. 5), 50 ug/ml cylohexamide (CHX) was added to inhibit protein synthesis. For live cell imaging of PIM-mCh-NLS cytosolic reaggregation protein synthesis was also inhibited using CHX or by washing out doxycycline 24 hours before imaging. In the latter condition, incubation with doxycycline was performed for 24 hours prior to wash out.

In order to prevent aggregate removal from the nucleus via mitosis during long term experiments (Fig. 1, 2), cell cycle arrest was performed by the addition of 2 mM thymidine.

### Live-cell imaging

#### Spinning Disc confocal microscopy, FRAP and photoactivation

Long term, high resolution, live-cell imaging of PIM-mCh-NLS removal from the nucleus was performed on a Nikon Ti2 microscope with a Perfect Focus System (PSF), a CSU-W1-T1 (Yokogawa) unit and a 100x oil emersion objective (Plan Apo λD, NA 1.45, Nikon). Cells were imaged in a TokaiHit incubation chamber (STXG-PLAMX-SETZ21L) on a MS-2000-XYZ stage with Piezo Top Plate (ASI). Illumination was performed using a 561 nm laser (Coherent Obis, Coherent) and images were captured on a sCMOS camera (Prime BSI). An z-stack (z-range: 5 μm, z-spacing: 0.5 μm) was captured every 10 seconds for 4 hours. The microscope was controlled by MetaMorph 7.8 (Molecular Devices).

Live-cell imaging of dualPIM removal from the nucleus following rapalog2 addition was performed on a Niken Eclipse Ti Microscope. The microscope was equipped with a CSU-X1-A1 confocal unit (Yokagawa), PFS, and a 60x oil immersion objective (Plan Apo VC, N.A. 1.40, Nikon). Cells were imaged on a motorized stage (MS-2000-XYZ, ASI) with a Piezo top plate and incubation chamber (INUBG2E-ZILCS, Tokai Hit). Illumination was performed using a 561 or 642 nm laser (Cobolt Jive and Voltrans Stradus, respectively). An Evolve 512 EMCCD camera (Photometrics) was used to capture images. Cells expressing dualPIM were imaged for 2 hours and every 1 minute a z-stack was captured (z-range: 9.5 μm, z-spacing: 0.5 μm). Cells were imaged for 2 hours and every 1 minute a z-stack was captured (z-range: 9.5 μm, z-spacing: 0.5 μm). Metamorph 7.7 software (Molecular Devises) was used to control the microscope.

Live-cell imaging of FRAP experiments following rapalog2 addition were performed on a Nikon Eclipse Ti2 Microscope. The microscope was equipped with a CSU-W1-A1 confocal unit (Yokagawa), PFS, and a 60x oil immersion objective (Plan Apo λD, N.A. 1.42, Nikon). Cells were imaged on a motorized stage (MS-2000-XYZ, ASI) with a Piezo top plate and incubation chamber (INUBG2E-ZILCS, Tokai Hit). Illumination was performed using a 561 nm laser (Coherent OBIS). A Prime BSI sCMOS camera (Photometrics) was used to capture images. A z-stack was captured every 45 seconds for 90 minutes (z-range: 5 μm, z-spacing: 1 μm). After 90 seconds, 2-3 aggregates per nucleus were bleached using an ILAS system (Gataca Systems). Bleaching was performed using a 561 nm laser (Coherent OBIS). Metamorph 7.10 software (Molecular Devises) was used to control the microscope.

FRAP experiments probing nucleocytoplasmic shuttling dynamics of PIM-mCh-NLS without rapalog2 addition and experiments using U2OS WT cells expressing PIM-mCh-PAGFP-NLS were done on asimilar confocal system to the one described above, equipped with a EMCCD camera (Photometrics Evolve 512). For the FRAP experiment cells were imaged every 30 seconds for 33 minutes (z-range: 5 μm, z-spacing: 1 μm). After 1 minute, a ROI placed outside the cell was bleached after every frame for 14 minutes. Subsequently, another ROI, placed in the cytosol at similar distance to the nucleus, was selected and bleached after every frame for 14 minutes. Lastly, a nuclear ROI was bleached for 4 minutes. U2OS WT cells expressing PIM-mCh-PAGFP-NLS were imaged every 2 minutes for 1.5-2 hours (z-range: 4 μm, z-spacing: 0.5 μm). After 2 minutes, 5 ROIs that did not contain nuclear PIM aggregates (based on mCherry signal) were photoactivated using a 405 nm laser (Vortran Stradus). After 20 minutes, 5 ROIs containing nuclear PIM aggregates were photoactivated. PAGFP was excited using a 491 nm laser (Vortran Stradus). Metamorph 7.7 software (Molecular Devises) was used to control the microscope.

Live-cell imaging of VCP (and VCP cofactor) overexpression was performed on a Nikon Eclipse Ti2 Microscope with a perfect focus system (PFS) and a 40x oil immersion objective (CFI Plan Fluor, NA 1.3, Nikon). Cells were imaged in a TokaiHit incubation chamber (STXG-WSKMX-SET). A CoolLED pe4000 (CoolLED) LED device was used for illumination in combination with ET-mCheery (49008, Chroma) and ET-Narrow-Band EGFP (49020, Chroma) filters. A z-stack was acquired (z-range: 1.5 μm, z-spacing 0.75 μm). GFP signal was used to determine the nuclear outline during analysis. Images were captured every 10 minute for 5 hours after rapalog2 removal. All images were acquired with an Andor SONA cMOS camera. The microscope was controlled using μManager software (Edelstein et al., 2014).

All other live-cell imaging was performed on a Nikon Eclipse Ti equipped with an incubator chamber (Tokai Hit; INUG2-ZILCS0H2) on a motorized stage (ASI). A CoolLED pE4000 (CoolLED) LED device was used for illumination in combination with a ET-mCherry (49008, Chroma) filter. Cells were images using a 40x oil immersion objective (Plan Fluor, NA 1.3, Nikon). Differential interference contrast (DIC) microscopy was used to determine the nuclear outline during analysis. Images were captured every 5-15 minutes for 3-7 hours after rapalog2 addition. All images were acquired with a Coolsnap HQ2 CCD camera (Photometrics). The microscope was controlled using µManager software (Edelstein et al., 2014).

### Fixed cell imaging

Imaging of fixed cells after antibody labelling was done using an Axio Observer 7 SP with Definite Focus 3 (Zeiss) and a 40x water immersion objective (C-Apochromat, NA 1.20, Zeiss). A z-stack was acquired (z-range: 5 μm, z-spacing: 0.5 μm) and captured using an LSM980 Airyscan 2 detector in Multiplex CO-8Y sampling mode. The microscope was controlled by ZEN Blue v3.9 software. Images of siRNA knock-down experiments were acquired on a ZEISS Cell discoverer 7

### siRNA knockdown screen

For the siRNA knockdown screen RPE-PIM-mCh-NLS cells were seeded in a glass bottom 96-well plate and incubated with doxycycline. Directly after seeding, cells were transfected with siRNAs using RNAiMAX (Thermo Fischer) and various siRNAs. Per well 0.1 μl RNAiMAX and 0.1 μl siRNA (stock concentration 20 μM) were added to 20 μl Optimem (Gibco). This reaction mix was thoroughly mixed, incubated for 10 minutes and subsequently added to the cells (final concentration 20 nM, final dose 2 pmol siRNA per well). All biological replicates were done in duplo. ∼32 hours after transfection thymidine was added to the medium and doxycycline was replenished. The following day, doxycycline was washed out and the assay was started by drug inhibitions and rapalog2 addition as described previously. Aggregation was started by addition of rapalog2 for 1 hour. Induction of aggregation was timed so that the 96-well plate could be fixed at a single timepoint. Cells were washed 1X with 1X-PBS and fixed using 4% PFA in 1X-PBS. Cells were stained using DAPI and kept in 1X-PBS until imaging. The following si-RNAs were used in this study. Scrambled si-Ctrl (Qiagen, 1027281), si-PSMC2 (Smartpool; L-008180-01, Dharmacon), si-PSMC5 (Smartpool; L-009484-00-0005, Dharmacon), si-PSMB2 (Smartpool; L-010461-00-0005, Dharmacon), si-HSPA1A (Smartpool; L-005168-00-0005, Dharmacon), si-DNAJB1 (Smartpool; L-012735-01-0010, Dharmacon), si-DNAJB6 (custom designed; CAUUCCAACAAUCUCGUAA, Dharmacon), and si-VCP (Smartpool; L-008727-00-0005, Dharmacon).

### Western blot analysis of PIM degradation in the nucleus

For biochemical analysis of nuclear PIM-NLS levels, RPE-PIM-mCh-NLS cells were incubated with doxycycline and thymidine before rapalog2 addition (24 hours and 14 hours, respectively). 30 minutes prior to rapalog 2 addition, cells were incubated with CHX and leptomycin B alone, or CHX and leptomycin B with NMS-873 or MG132. Rapalog2 was added as described previously and cells were incubated for 6 hours. Cells were washed 1X with ice-cold 1X-PBS, lysed using Laemmli buffer and processed for western blot analysis.

### Immunofluorescence and Western blot labelling

The following primary antibodies were used in this study: rabbit anti-RFP (Rockland, ref: 600-401-379, lot: 46510), rabbit anti-LC3 (MBL, ref: PM036, lot: 037), mouse anti-alpha tubulin (Sigma Aldrich, ref: T-5168, lot: 0000093552, clone: B-5-1-2), rabbit anti-VCP (abcam, ref: ab111740), rabbit anti-HSP70 (ProteinTech, ref: 10995-1-AP), rabbit anti-DNAJB1 (ProteinTech, ref: 13174-1-AP), rabbit anti-DNAJB6 (gift from Braakman group), rabbit anti-PSMC2 (Atlas Antibodies, ref: HPA019238), rabbit anti-PSMC5 (Atlas Antibodies, ref: HPA064293), and rabbit anti-PSMB2 (ProteinTech, ref: 15154-1-AP). The following secondary antibodies were used in this study: goat anti-rabbit AF488 (Thermo Fischer Scientific, ref: A-11034, lot: 2541675), goat anti-mouse HRP (Dako, ref: PO447, lot: 41456532), swine anti-rabbit HRP (Dako, ref: PO399, lot: 20069482). Western blot membranes were developed using ECL Prime western blotting detection reagent (Sigma Aldrich, ref: GERPN2236).

### Quantifications

The clearance of single aggregates from the nucleus was analysed by manually drawing a ROI around an aggregate and measuring the mCherry signal intensity. The number of particles, total particle intensity and average particle intensity of PIM-NLS aggregates in the nuclei of single cells was analysed by manually tracing the nuclear outline of a cell (at 30-minute intervals). We then integrated an updated version of the ComDet plugin (Comdet v.0.5.5.; https://github.com/UU-cellbiology/ComDet), into a custom macro that sequentially detects particles with a standard deviation (SD) of 10, and an estimated size of 4 pixels. For analysis of nuclear PIM aggregation after si-RNA knockdown, z-stacks were first combined to produce a maximum projection. Then, Stardist 2D (https://github.com/stardist/stardist) was used to segment nuclear ROIs based on the DAPI signal. In the mCherry channel, these ROIs were used to determine the mean signal intensity per nucleus, and the SD of the mCherry signal of all pixels within one nucleus. The mean intensity and SD were then used to calculate the coefficient of variation for each nucleus (referred to as the aggregation coefficient in Fig. 2, D). In order to estimate effect sizes, we calculated a cutoff value based on the mean of the si-Control sample (mean + 2*SD).

The FRAP experiment probing nucleocytoplasmic shuttling dynamics was analysed by measuring the nuclear PIM-mCherry-NLS signal of an ROI that wat placed in the nucleus but did not overlap with any ROIs selected for photobleaching. Nuclear PIM-mCherry-NLS intensity was normalized to the first 3 frames of the acquisition, before onset of bleaching. The decay rate was estimated by a linear regression fit of the slopes during the bleaching of the ROI outside the cell and the cytosolic ROI. All other FRAP experiments were analysed by manually measuring single aggregate intensity values. Intensity values were background subtracted and normalized to the pre-bleaching aggregate intensity.

All quantifications were performed on raw, unprocessed images or sum projections of z-stacks. All analysis were done using FIJI (Schindelin et al., 2012).

### Predictive Analysis

Sequence-based prediction of nuclear export signals (NESs) within the PIM-mCh-NLS protein was performed with NESmapper (Kosugi et al., 2014), run using Strawberry Perl (5.42.0.1-64bit; https://strawberryperl.com/). Then, structure predictions for full-length PIM-mCh-NLS and the predicted NES sequences were performed with the AlphaFold 3 server (v3.01) (Abramson et al., 2024). Subsequent molecular graphics and surface-area calculations were performed with UCSF ChimeraX (1.8) (Meng et al., 2023). Finally, protein-protein docking simulations for full-length PIM-mCh-NLS to CRM1 and a predicted NES sequence to CMR1 were performed using the HADDOCK (High Ambiguity Driven protein-protein DOCKing) server (Honorato et al., 2021, 2024) (v2.5-2025.08). Crystal structures for free CRM1 and the CRM1-SNUPN complex were acquired from the RCSB Protein Data Bank (RCSB PDB) (PDB IDs: 4FGV, 3GB8) (RCSB.org) (Berman et al., 2000). Sequence-based prediction of nuclear import signals (NLSs) within the PIM-mCh-NLS protein was performed with the NLSExplorer server (Y. F. Li et al., 2025) and the NLStradamus server (Nguyen Ba et al., 2009) (2 state HMM static, 2 state HMM dynamic, and 4 state HMM static models were all used; prediction cutoff 0.5).

### Statistical analysis

All graphs were generated using Graphpad Prism (v.9.4.0). Graphad was also used to perfom statistical tests.

---

**Figure S1:**
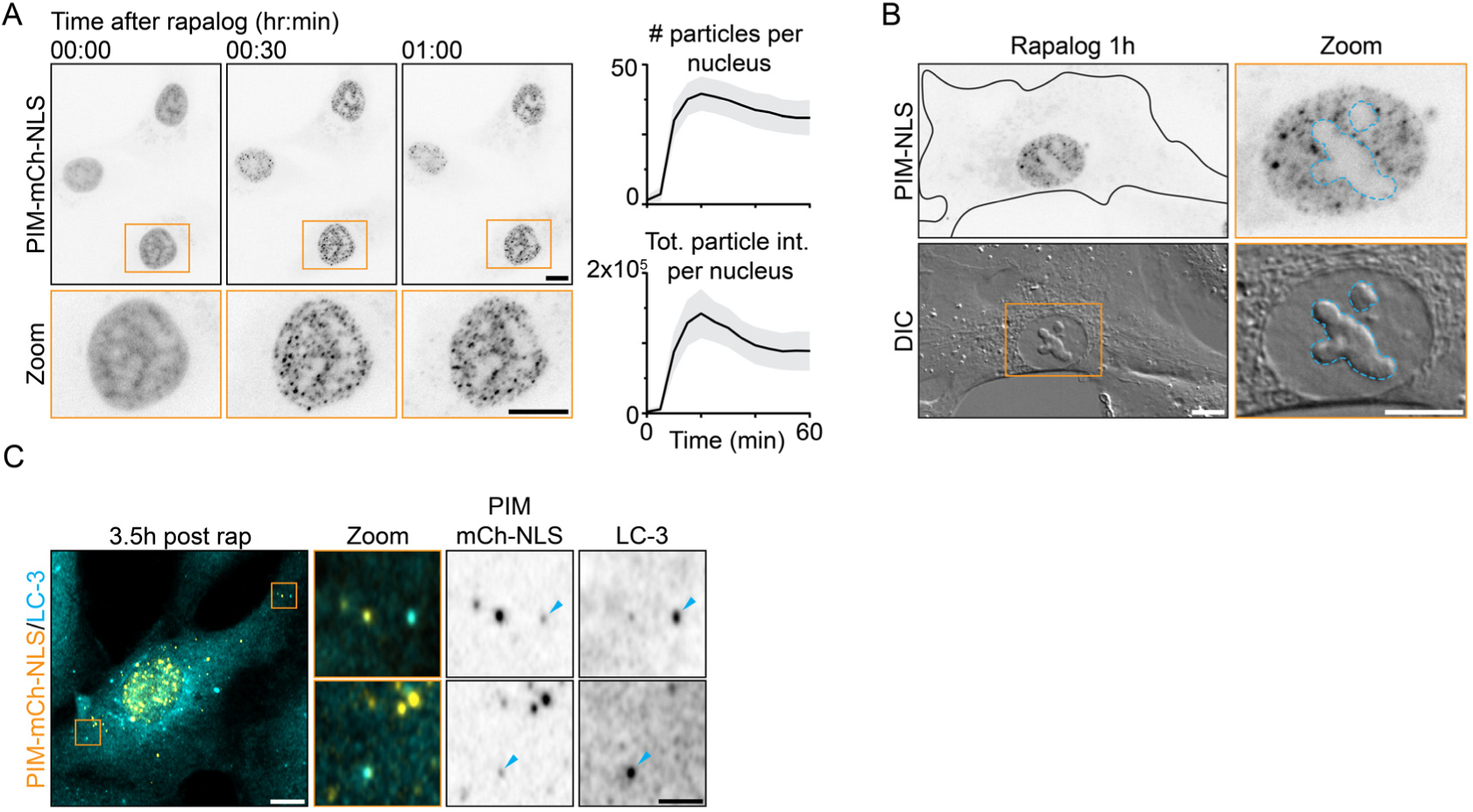
Formation and clearance of nuclear PIMs. (**A**) Representative live-cell images of RPE-PIM-mCh-NLS cells. Addition of rapalog after onset of imaging results in the rapid (∼1 hour) formation on nuclear aggregates. Zooms (orange boxes) show aggregate formation in a single nucleus. Traces indicate the mean and the shaded area represents 95% confidence interval. (**B**) Representative image of RPE-PIM-mCh-NLS cell 1 hour after rapalog addition. Zoom shows a single nucleus and DIC was used to visualize the nucleoli (outlined in cyan). (**C**) Representative z-projection of confocal images of fixed RPE-PIM-mCh-NLS cells (orange) 3.5 hours after rapalog addition. Cells were stained using LC-3 (cyan). Zooms show regions with cytosolic PIM-mCh-NLS aggregates that colocalize with LC-3 (cyan arrowheads). Scale bars represent 10 μm (A, B and C) or 2.5 μm (Zooms of C).

**Figure S2:**
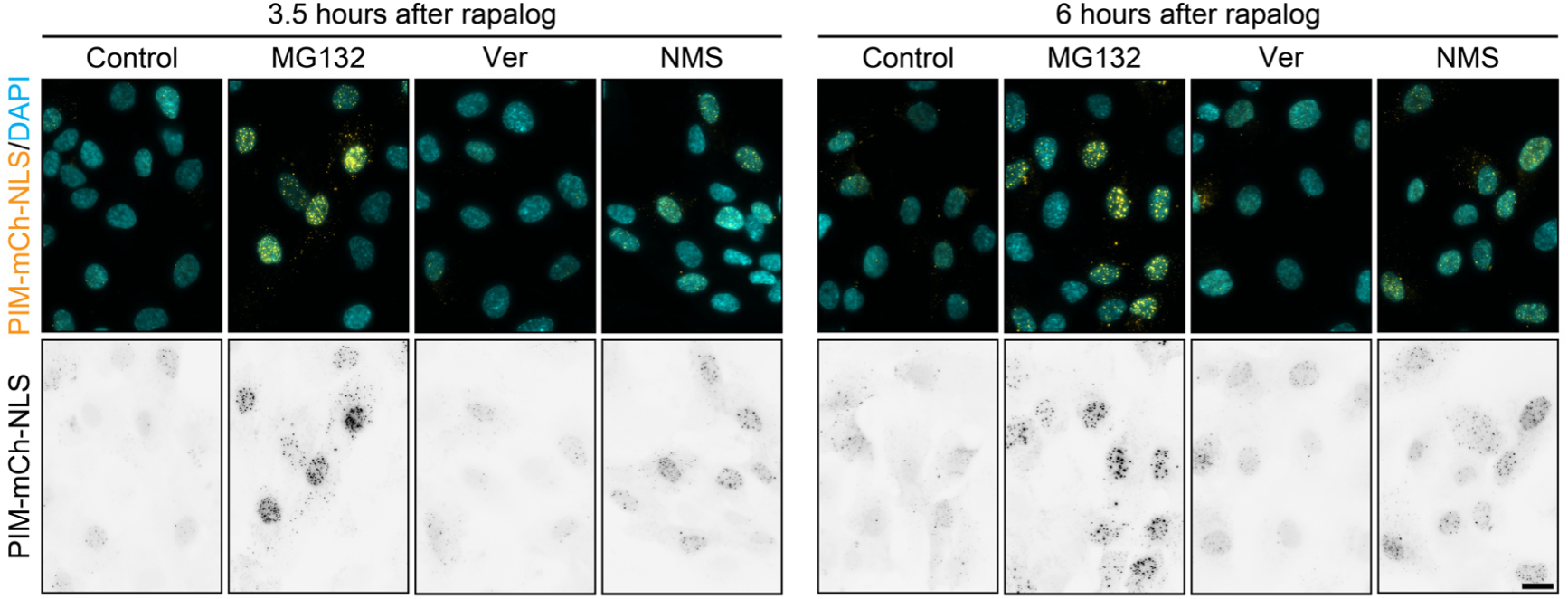
Inhibition of the proteasome, Hsp70 and VCP differentially affects nuclear PIM clearance. Representative images of RPE-PIM-mCh-NLS cells (orange or inverted gray scale) fixed 3.5 or 6 hours after rapalog induced PIM aggregation. Images show control cells, and cells treated with MG132 (proteasome inhibitor), Ver-155008 (Hsp70 inhibitor) and NMS-873 (VCP inhibitor). DAPI (cyan) was used to visualize nuclei. Scale bars represents 20 μm.

**Figure S3:**
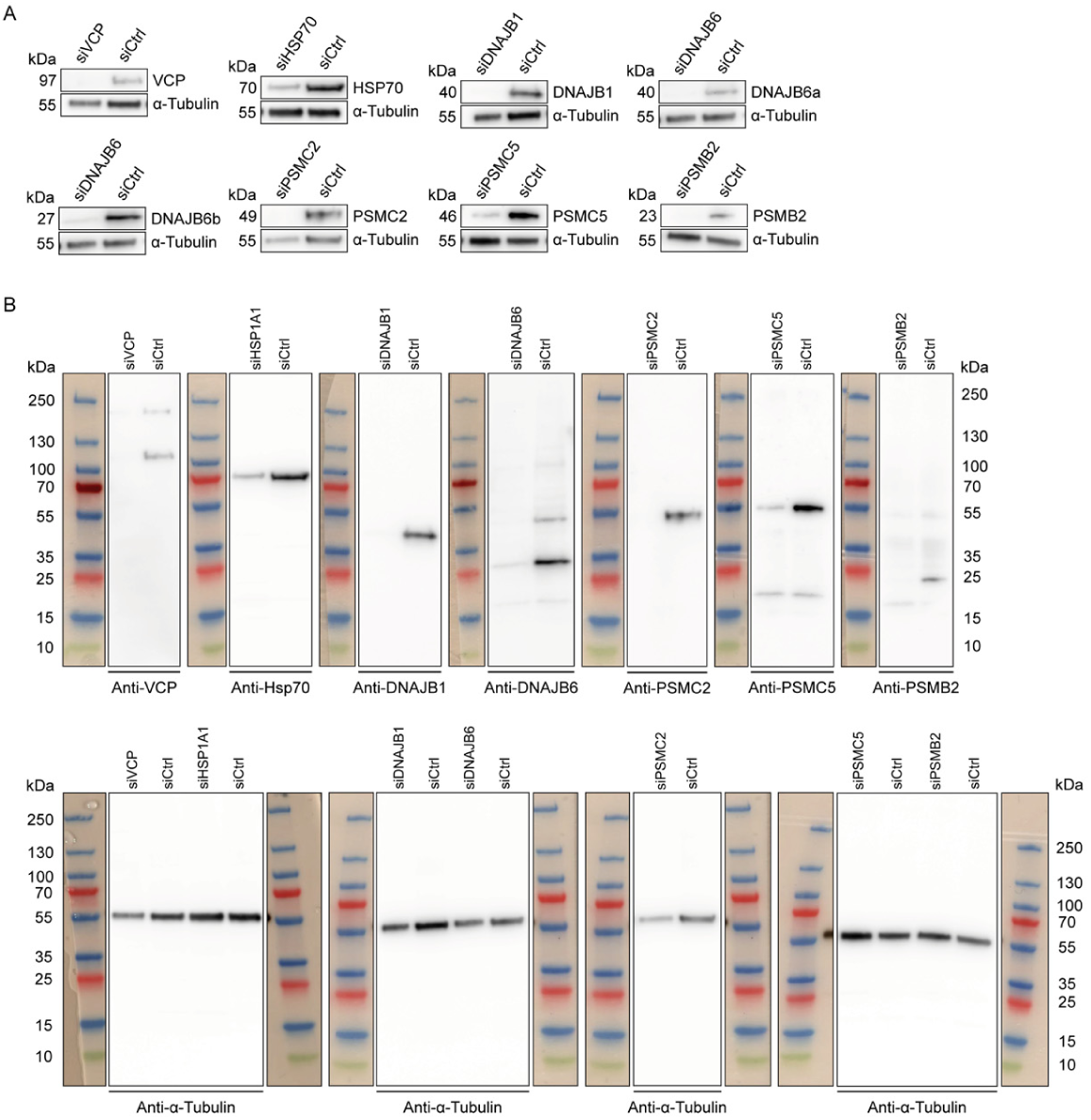
Western Blot analysis of siRNA knockdown panel. (A) Validation of knockdown efficiency for siRNAs used in figure 2. (B) Complete blots showing full protein range.

**Figure S4:**
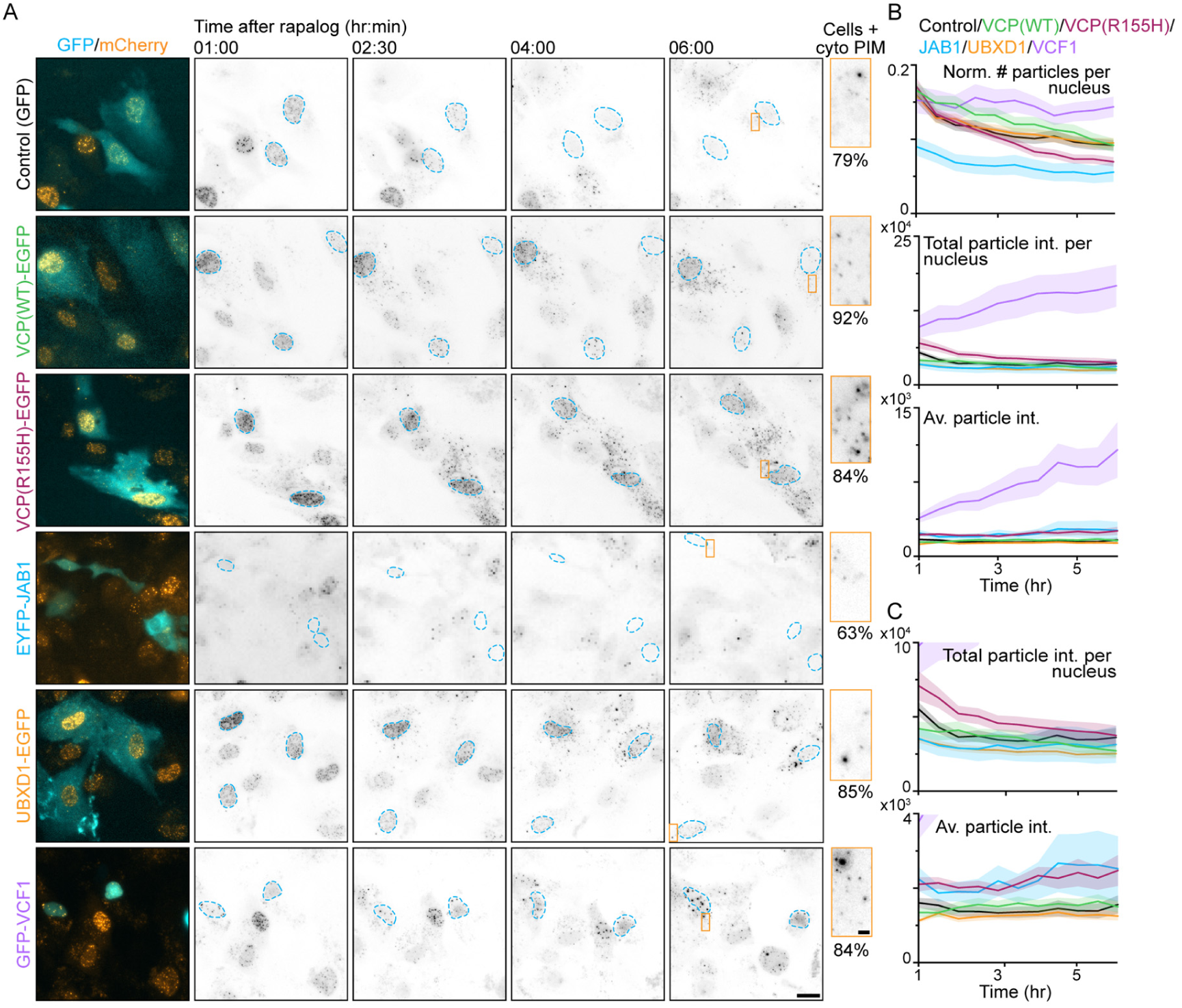
Overexpression of VCP and VCP cofactors. (A) Representative images of live-cell imaging of RPE-PIM-mCh-NLS cells transfected with GFP, VCP(WT)-EGFP, VCP(R155H)-EGFP, UBXD1-GFP, EYFP-JAB1, or GFP-VCF1. Orange boxes indicate a cytosolic region that is shown as a zoom, highlighting cytosolic aggregates. (B) Quantification showing the number of particles per nucleus (normalized to nucleus area) (top graph), total intensity of particles per nucleus (middle graph), or the average particle intensity per nucleus (bottom graph) over time. Traces represent the mean and shaded areas show the SEM of GFP (grey), VCP(WT) (green), VCP(R155H) (magenta), JAB1 (cyan), UBXD1 (orange), and VCF1 (purple) overexpressing cells (N≥3, n_cells_ = 104, 64, 100, 146, 43, 37, respectively). (C) Particle intensity graphs as shown in B with a reduced y-axis. Nuclei are outlined in cyan. Scale bars represent 20 μm and 2 μm (zoom).

**Figure S5:**
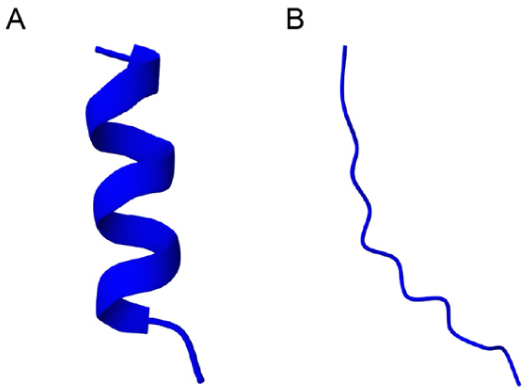
Folded structure of isolated NES motifs. (A) AlphaFold3-predicted structure for NES1 sequence. (B) AlphaFold3-predicted structure for NES2 and NES2B sequences.

**Figure S6:**
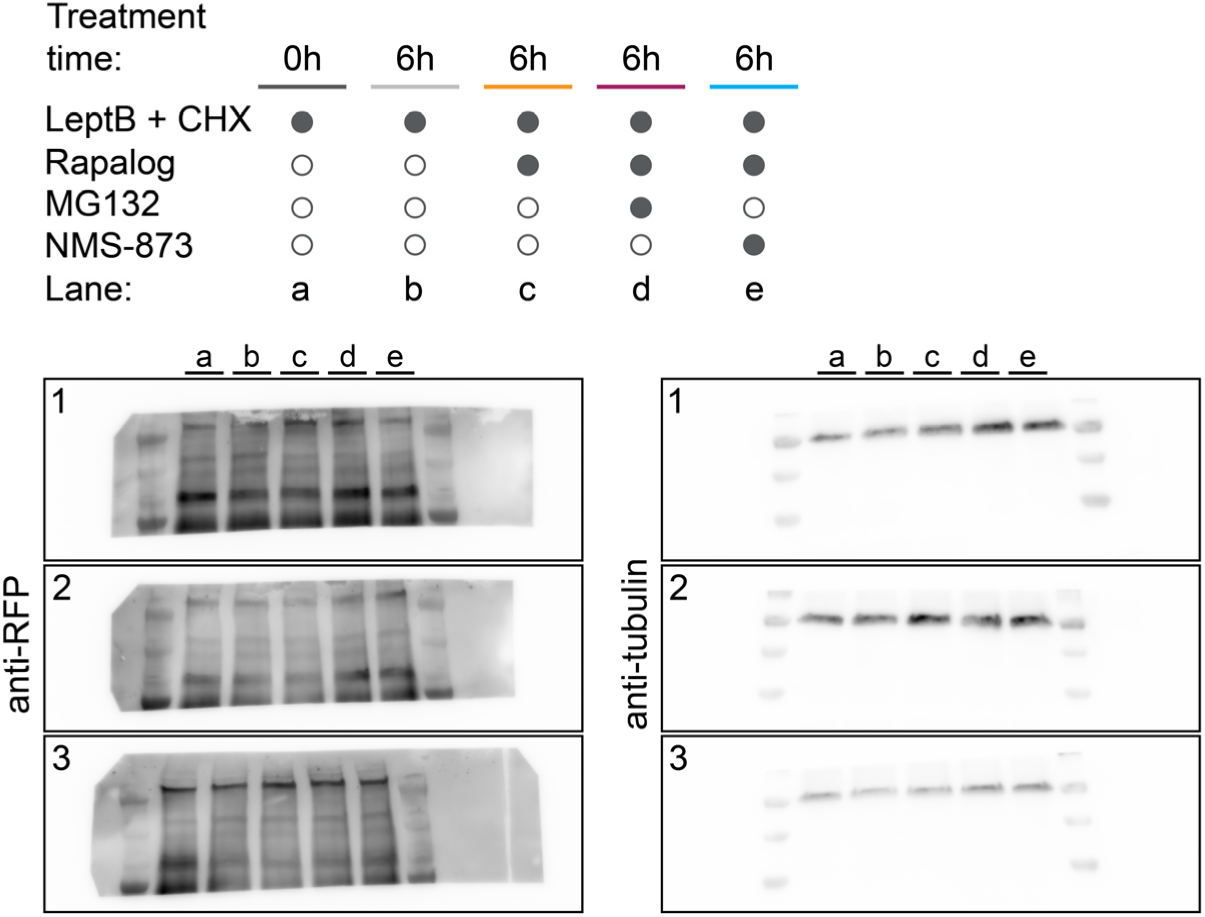
Western blot analysis of PIM-mCherry-NLS levels. Complete and unspliced images of blots used in figure 4 (D). Membranes (1-3) were cut and subsequently stained using anti-RFP or anti-alpha tubulin.

**Table S1:** NLS predictions for PIM-mCh-NLS.

|  |  |
| --- | --- |
| Model | Predicted NLS sequence |
| NLSExplorer | SEFPKKKRKV |
| NLStradamus 2 state HMM static | KKKRKV |
| NLStradamus 2 state HMM dynamic | KKKRKV |
| NLStradamus 4 state HMM static | PKKKRKV |

**Table S2:** NESmapper output for PIM-mCh-NLS.

| Position | Sequence | Score | NES Class |
| --- | --- | --- | --- |
| 60 | IRGWEEGVAQMSVG | 16.2 | 1a |
| 98 | HATLVFDVELLKLE | 16.2 | 1a |
| 169 | IRGWEEGVAQMSVG | 16.2 | 1a |
| 207 | HTTLVFDVELLKLE | 11.34 | 1a |
| 522 | IRGWEEGVAQMSVG | 16.2 | 1a |
| 560 | HATLVFDVELLKLE | 16.2 | 1a |
| 631 | IRGWEEGVAQMSVG | 16.2 | 1a |
| 669 | HATLVFDVELLKLE | 16.2 | 1a |
| 747 | IRGWEEGVAQMSVG | 16.2 | 1a |
| 785 | HATLVFDVELLKLE | 16.2 | 1a |
| 856 | IRGWEEGVAQMSVG | 16.2 | 1a |
| 894 | HATLVFDVELLKLE | 16.2 | 1a |

